# Ulk1(S555) inhibition alters nutrient stress response by prioritizing amino acid metabolism

**DOI:** 10.1101/2025.06.30.662412

**Authors:** Orion S. Willoughby, Anna S. Nichenko, Matthew H. Brisendine, Niloufar Amiri, Shelby N. Henry, Daniel S. Braxton, John R. Brown, Braeden J. Kraft, Kalyn S. Jenkins, Adele K. Addington, Alexey V. Zaitsev, Steven T. Burrows, Ryan P. McMillan, Haiyan Zhang, Spencer A. Tye, Charles P. Najt, Siobhan E. Craige, Timothy W. Rhoads, Junco S. Warren, Joshua C. Drake

## Abstract

Metabolic flexibility, the capacity to adapt fuel utilization in response to nutrient availability, is essential for maintaining energy homeostasis and preventing metabolic disease. Here, we investigate the role of Ulk1 phosphorylation at serine 555 (S555), a site regulated by AMPK, in coordinating metabolic switching following short-term caloric restriction and fasting. Using Ulk1(S555A) global knock-in mice, we show loss of S555 phosphorylation impairs glucose oxidation in skeletal muscle and liver during short-term CR, despite improved glucose tolerance. Metabolomic, transcriptomic, and mitochondrial respiration analyses suggest a compensatory reliance on autophagy-derived amino acids in Ulk1(S555A) mice. These findings suggest Ulk1(S555) phosphorylation as a critical regulatory event linking nutrient stress to substrate switching. This work highlights an underappreciated role of Ulk1 in maintaining metabolic flexibility, with implications for metabolic dysfunction.

**Graphical Abstract:** 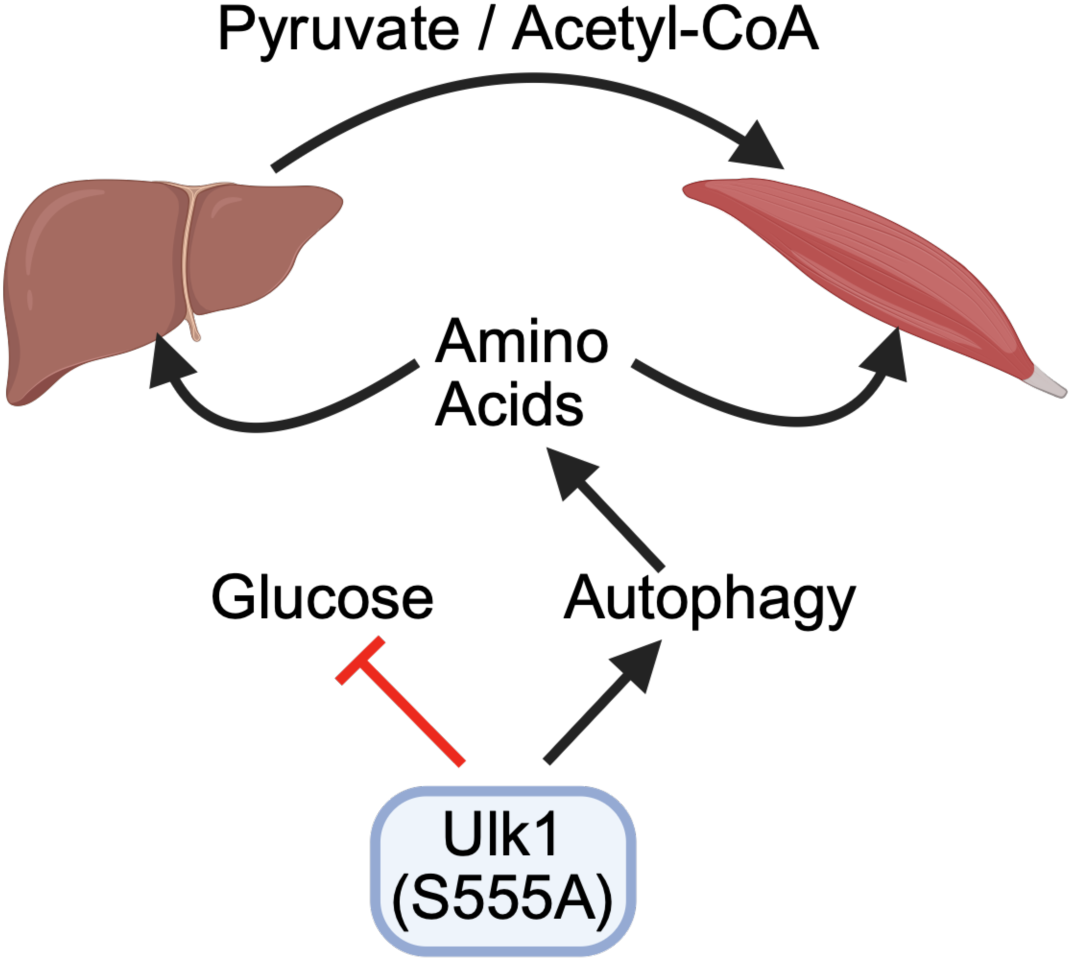

## Introduction

Metabolic flexibility, the ability to respond and adapt to energetic demands (Goodpaster & Sparks, 2017), is necessary for survival. Regularly inducing a metabolic switch, such as switching from glucose to fatty acid oxidation during regular intervals of fasting, as occurs during intermittent fasting (IF) or caloric restriction (CR) has been shown to extend healthspan and lifespan in numerous organisms (Duregon et al., 2021; Lee et al., 2021). Consequently, metabolic inflexibility indicates metabolic dysfunction, contributing to many metabolic-related diseases (e.g., diabetes, sarcopenia, and obesity) and characteristic of aging (Riera & Dillin, 2015; Zhou et al., 2023). Skeletal muscle is the primary utilizer of glucose, accounting for 60-80% of glucose metabolism (Ng et al., 2012). CR and/or IF increase capacity to take up glucose (Pak et al., 2021; Teong et al., 2023), particularly in skeletal muscle (Sharma et al., 2014; Wang et al., 2017), as well as mitochondrial enzymatic processes (e.g., tricarboxylic acid cycle and oxidative phosphorylation) to utilize the pyruvate produced from the breakdown of glucose through glycolysis (Chen et al., 2015; Pak et al., 2021; Rhoads et al., 2020). Depletion of glucose with fasting initiates gluconeogenesis, with skeletal muscle providing amino acids via protein breakdown to be transaminated to pyruvate primarily in the liver via the Cahill cycle (Petersen et al., 2019) or within the muscle itself (Neinast et al., 2019) to meet skeletal muscle energy requirements. However, the utilization of amino acids as glucogenic precursors is inefficient, with 20-33% of the energy from amino acid-derived pyruvate being lost to the energetic cost of protein breakdown and urea synthesis (Hall, 2006; Veldhorst et al., 2009).

A primary axis for the molecular response to changes in cellular energy is coordination between 5’-AMP-activated kinase (AMPK) and mechanistic target of rapamycin 1 (mTORC1) (Green et al., 2022). AMPK responds to increased AMP/ADP by suppressing energetically costly processes, like protein synthesis, to conserve cellular energy, while mTORC1 promotes anabolism during energetic abundance. A shared substrate between AMPK and mTORC1 is Unc-51-like autophagy activating kinase (Ulk1), which is best characterized for its role in recycling cellular components through autophagy in response to energetic stress (Pareek & Kundu, 2024). Inducible loss of Ulk1 in skeletal muscle is sufficient to impair exercise training-mediated improvements in glucose tolerance (Drake et al., 2021; Laker et al., 2017), whereas silencing downstream autophagy genes (e.g., *ATG5, ATG7, BCN1*) *in vitro* does not affect glucose metabolism (Li et al., 2016), suggesting a direct role for Ulk1 in metabolic regulation. Indeed, *in vitro,* Ulk1 has been shown to directly phosphorylate glycolytic enzymes, hexokinase (HK), phosphofructokinase 1 (PFK1), and enolase 1 (ENO1), as well as the gluconeogenic enzyme fructose-1,6-bisphosphatase (FBP1) in response to nutrient stress (Li et al., 2016). Ulk1-dependent glucose metabolism may function to modulate protein synthesis only when sufficient energy is available to support the energetic investment (Park et al., 2023; Yoon et al., 2020). Despite this evidence for Ulk1 having regulatory functions modulating metabolic responses to energetic stress, phosphorylation site-dependent roles for Ulk1 in modulating metabolism *in vivo* are not well defined.

AMPK phosphorylates Ulk1 at multiple sites (Park et al., 2023), including S467, S555, T574, and S637 (murine sites) (D. F. Egan et al., 2011). Of these, S555 is most readily phosphorylated in response to energetic stressors like exercise (Laker et al., 2017) and fasting (Bujak et al., 2015) *in vivo*. Based on evidence that deletion of Ulk1 or simultaneous impairment of S467, S555, T574, and S637 (4SA) disrupts the autophagy response to energetic stress (D. F. Egan et al., 2011; Laker et al., 2017), we and others have interpreted the phosphorylation of Ulk1 at S555 to initiate its autophagy function, as S555 is most responsive *in vivo* and precedes autolysosome formation (Bujak et al., 2015; D. F. Egan et al., 2011; Laker et al., 2017). However, recent evidence in cell culture models has challenged this paradigm by demonstrating that AMPK-dependent Ulk1(S555) phosphorylation, in coordination with other AMPK-dependent phosphorylation events, inhibits autophagy to conserve cellular energy when glucose is limited, depending on the availability of amino acids (Kazyken et al., 2024; Park et al., 2023). CRISPR-engineered Ulk1(S555A) mice maintain a higher RER during exercise, delayed crossover time (RER < 0.85), and have elevated circulating lactate following exercise in the absence of an effect on autophagy/mitophagy (Guan, Spaulding, et al., 2024). Therefore, phosphorylation of Ulk1 at S555 may have an unappreciated role in mediating metabolic flexibility *in vivo*.

Here, we utilized Ulk1(S555A) global knock-in mice (Guan, Spaulding, et al., 2024) to interrogate site-specific requirement(s) of Ulk1(S555) phosphorylation for metabolic flexibility under different durations of nutrient stress. We discovered inhibition of Ulk1(S555) altered the rate of body weight loss during 8 weeks of 40% caloric restriction (CR) and blunted CR-mediated increases in glucose oxidation capacity of skeletal muscle and liver via U-[^14^C]-glucose tracing, despite improved glucose tolerance. GC-MS-based metabolomics, as well as bulk RNA sequencing of skeletal muscle and liver, suggests increased reliance on amino acids as energetic substrates during CR. Ulk1(S555A) skeletal muscle mitochondrial oxygen consumption is elevated in the presence of amino acids and continues to increase in response to glutamate/malate stimulation of Complex I respiration, yet fails to increase oxygen consumption in response to ADP stimulation of Complex V. In response to fasting, we confirmed Ulk1(S555A) mice have a delayed metabolic switch and maintain circulating urea that can be attenuated with the autophagy inhibitor 3-methyladenine (3-MA). These results demonstrate phosphorylation of Ulk1 at S555 is required for metabolic flexibility and suggest that loss of S555 biases metabolism towards utilization of amino acids through sustained autophagy.

## Results

### Attenuated glucose oxidative capacity following caloric restriction in the absence of Ulk1(S555) phosphorylation

The effects of caloric restriction (CR) are associated with extended healthspan and lifespan and amelioration of age-related pathologies, in part, through improved coordination between skeletal muscle and liver metabolism, leading to improved glucose tolerance and homeostasis, insulin sensitivity, and metabolic flexibility (Fok et al., 2013; Matyi et al., 2018). WT and Ulk1(S555A) mice were designated to either *ad libitum* (AL) or 8-week 40% CR (Fig. 1A). Food intake per cage for CR was determined by measuring three consecutive days of food consumption (Fig. 1A; S1A). AL Ulk1(S555A) mice maintained higher body mass over the course of the study compared to WT mice (Fig.1B). CR groups decreased body mass, with Ulk1(S555A) mice demonstrating a diminished response in body mass loss to CR compared to CR WT mice (Fig. 1B-C). WT mice maintained lean and fat mass as a percentage of body mass over the duration of CR (Fig. S1B, C). Ulk1(S555A) mice, however, maintained significantly more fat mass (Fig. S1B, C). Pre- and post-CR glucose tolerance tests (Fig. 1D) demonstrated improved glucose clearance in both WT and Ulk1(S555A) with CR (Fig. 1E-F), although Ulk1(S555A) were overall less glucose tolerant than WT (Fig. 1E). Contrary to deletion of Ulk1, which blunts exercise training-mediated improvements in glucose tolerance and insulin sensitivity (Drake et al., 2021; Laker et al., 2017), our findings suggest inhibition of S555 does not overtly affect improvement in glucose tolerance with CR, which is analogous to our recent findings following exercise training (Guan, Spaulding, et al., 2024).

**Figure 1.**
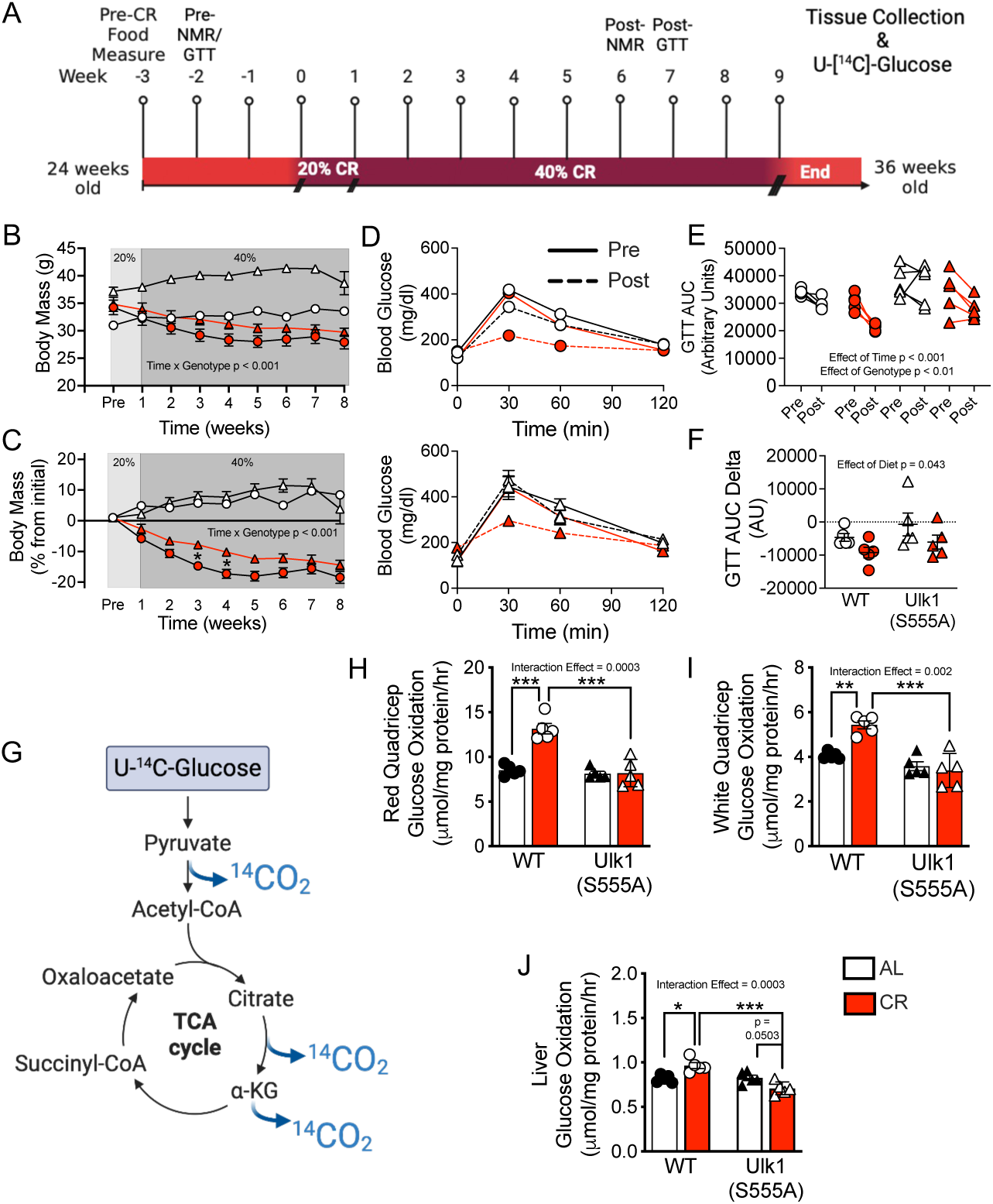
Attenuated glucose oxidative capacity following caloric restriction in the absence of Ulk1(S555). A) Caloric restriction study design. B) Weekly body mass (g) during CR. C) Weekly body mass % change from baseline during CR. D) glucose tolerance test. E) GTT area under the curve. F) GTT AUC delta. G) Uniformly labeled U-^14^C-Glucose tracer metabolic pathway. H) U-^14^C-Glucose oxidation in red fibers isolated from quadriceps. I) U-^14^C-Glucose oxidation in white fibers isolated from quadriceps. J) U-^14^C-Glucose oxidation in liver. n=5 per group. Data presented as mean ± SEM and analyzed via repeated measures or two-way ANOVA with post-hoc analysis when a significant interaction effect was found (* = p <0.05, ** p < 0.01, *** p < 0.0001).

Skeletal muscle is the primary site for glucose metabolism during nutrient stress (Ng et al., 2012). To assess if increased glucose tolerance with CR was accompanied by an increased capacity to oxidize glucose, we utilized U-[^14^C]-Glucose to uniformly assess oxidation of all carbons within glucose through glycolysis and the TCA cycle (Fig. 1G) (Specht et al., 2021). CR increased U-[^14^C]-Glucose oxidation in both red (oxidative) and white (glycolytic) muscle fibers from quadricep, as well as liver of WT mice (Fig. 1H-J), suggesting an increased capacity to oxidize glucose following short-term CR. However, glucose oxidation capacity, as assessed by U-[^14^C]-Glucose, did not change in skeletal muscle or liver from Ulk1(S555A) mice (Fig. 1H-J). This suggests phosphorylation of Ulk1 at S555 (Fig. S1D-F) may have undefined roles in metabolic regulation to meet skeletal muscle and liver energetic needs during prolonged nutrient stress, with loss of Ulk1(S555) causing an irregular shift away from glucose-derived substrates.

### Caloric restriction promotes skeletal muscle amino acid metabolism in Ulk1(S555A) mice

As Ulk1(S555A) mice did not increase capacity to oxidize glucose in skeletal muscle following 8 weeks of 40% CR, we performed GC-MS-based metabolomics and bulk RNAseq of skeletal muscle (quadricep) to understand how loss of Ulk1(S555) alters metabolic adaptation to CR. Metabolic pathway impact of skeletal muscle from Ulk1(S555A) mice following CR showed eight of the top 10 upregulated pathways were related to amino acid metabolism (Fig. 2A-B; S2B). Analysis of individual amino acids showed skeletal muscle from Ulk1(S555A) mice had elevated levels of glucogenic and ketogenic amino acids that were either elevated in Ulk1(S555A) and unaffected by CR (e.g. glycine, methionine, and lysine, with alanine and threonine nearing significance) or further elevated in response to CR (e.g. serine) (Fig. S2A-C). Serine is a principal donor in one-carbon units to the folate cycle and serves as a carbon source and redox regulator, and increased serine catabolism has been shown to suppress glucose oxidation (Yang et al., 2020). Other amino acids, such as alanine, are well defined as being utilized as energetic substrates within skeletal muscle (Felig, 1973; Hewton et al., 2021). Consistent with the metabolic pathway analysis (Fig. 2B), RNA sequencing from skeletal muscle of Ulk1(S555A) mice showed an increased enrichment score for nitrogen metabolism and biosynthesis of amino acids in AL mice (Fig. 2C) and arginine and proline metabolism with CR (Fig. 2D). Metabolomics of skeletal muscle showed elevated α-ketoglutarate in response to CR in Ulk1(S555A) mice coinciding with elevated serine and alanine (Fig. 2E-G). This is consistent with the transfer of the amino group from glutamate to pyruvate to generate alanine and/or serine and α-ketoglutarate (Fig. 2E) and positive correlation with amino acid biosynthesis (Fig. 2C). Further, Ulk1(S555A) mice had significantly less citric acid in skeletal muscle and less expression of *Pdp2* and *Pik3r1* (Fig. S2D), collectively supporting impaired glucose oxidation and improved tolerance in skeletal muscle (Fig. 1H, I). These data suggest impaired glucose oxidative capacity in skeletal muscle following CR in the absence of Ulk1(S5555) may result in glucose being diverted towards amino acid synthesis as an alternative energy source.

**Figure 2.**
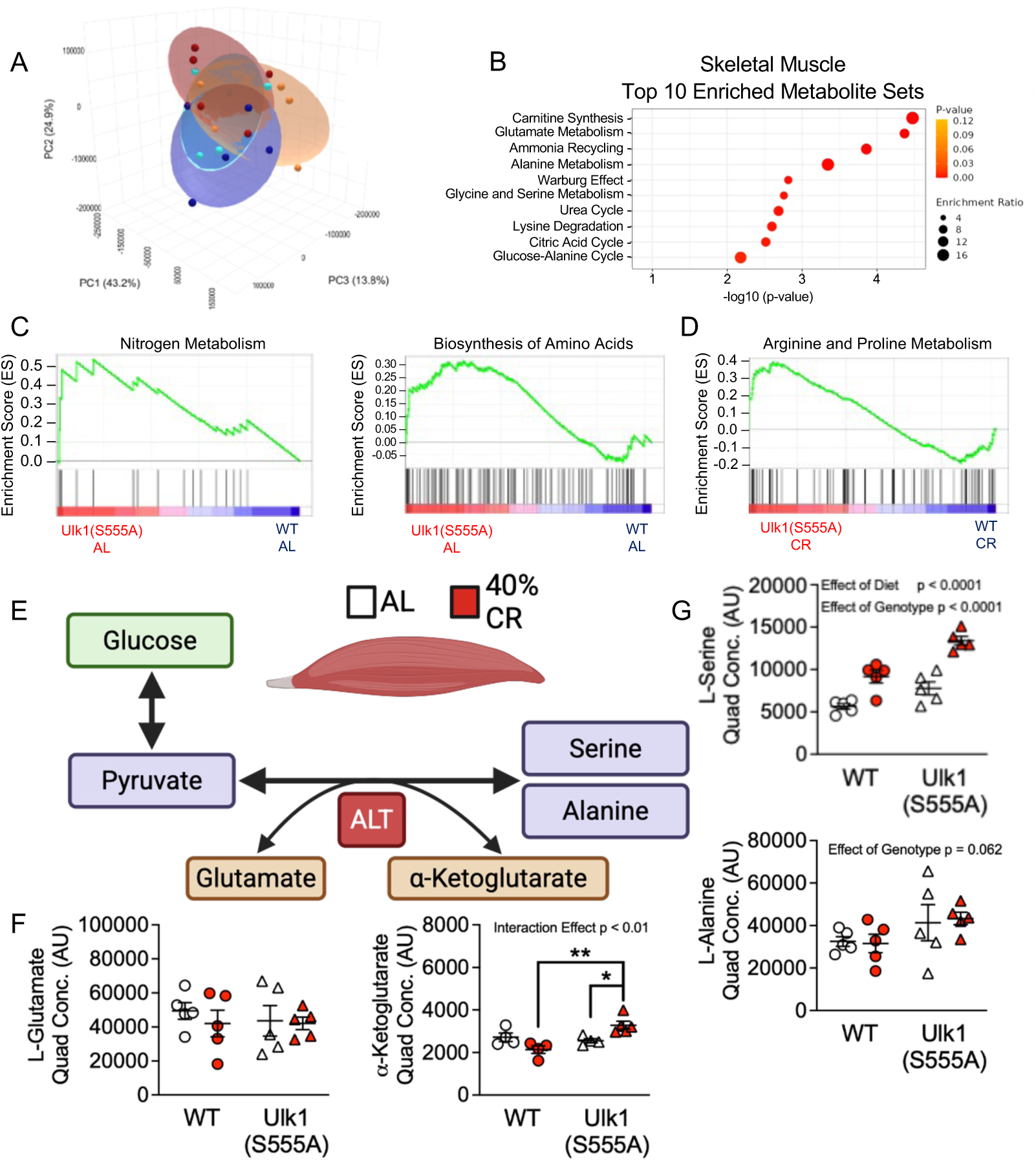
Caloric restriction leads to possible reliance on amino acids in skeletal muscle in absence of Ulk1(S555). A) PCA plot of skeletal muscle metabolites. B) Enriched metabolites of skeletal muscle. C) Skeletal muscle enrichment scores of nitrogen metabolism and amino acid biosynthesis in WT and Ulk1 AL groups from bulk RNA sequencing. D) Enrichment score of arginine and proline metabolism in WT and Ulk1 CR groups from bulk RNA sequencing. E-G) Alanine and Serine biosynthesis pathway and metabolite analysis. n=5 per group. Data presented as mean ± SEM and analyzed via two-way ANOVA with post-hoc analysis when a significant interaction effect was found (* = p < 0.05, ** = p < 0.01).

### Loss of Ulk1(S555) phosphorylation increases skeletal muscle mitochondrial oxidative capacity of amino acids

Alanine and serine can be metabolized either in the cytoplasm or imported and transaminated in mitochondria to pyruvate to supply the TCA cycle or into the reducing equivalent NADH (serine) for the electron transport chain (Fig.3A) (Lucas et al., 2018; Parker et al., 2020; Wu et al., 2023; Yang et al., 2020). To determine if alanine and/or serine in skeletal muscle of Ulk1(S555A) mice (Fig.2G) reflected an increased capacity to utilize amino acids as energetic substrates, we assessed respiration of mitochondria isolated from quadricep skeletal muscle. WT and Ulk1(S555A) mice were fasted overnight (∼12 hours) and mitochondria were isolated and incubated in the presence of increasing concentrations (2mM, 5mM, 10mM) of either alanine or serine. Baseline oxygen consumption rate (OCR) was significantly elevated in the presence of alanine in a concentration-dependent manner in mitochondria isolated from Ulk1(S555A) skeletal muscle (Fig.3B-C). Baseline OCR was also elevated in Ulk1(S555A) mitochondria compared to WT in the presence of serine at all concentrations (Fig.3F-G). In the presence of alanine or serine, OCR remained elevated in Ulk1(S555A) mitochondria compared to WT with the addition of glutamate/malate (10mM/5mM) to stimulate Complex I-dependent respiration, though serine caused a concentration-dependent suppression of OCR in both genotypes (Fig.3D, H). However, with the addition of ADP, which mimics energetic stress conditions, only WT mitochondria were able to further elevate OCR in the presence of alanine or serine at any concentration (Fig.3E, I). These data support the notion that skeletal muscle mitochondria have an increased capacity to utilize amino acids in the absence of Ulk1(S555) but that there is a lower limit on maximal mitochondrial energy production that may limit energy production during prolonged energetic stress, such as during CR.

**Figure 3.**
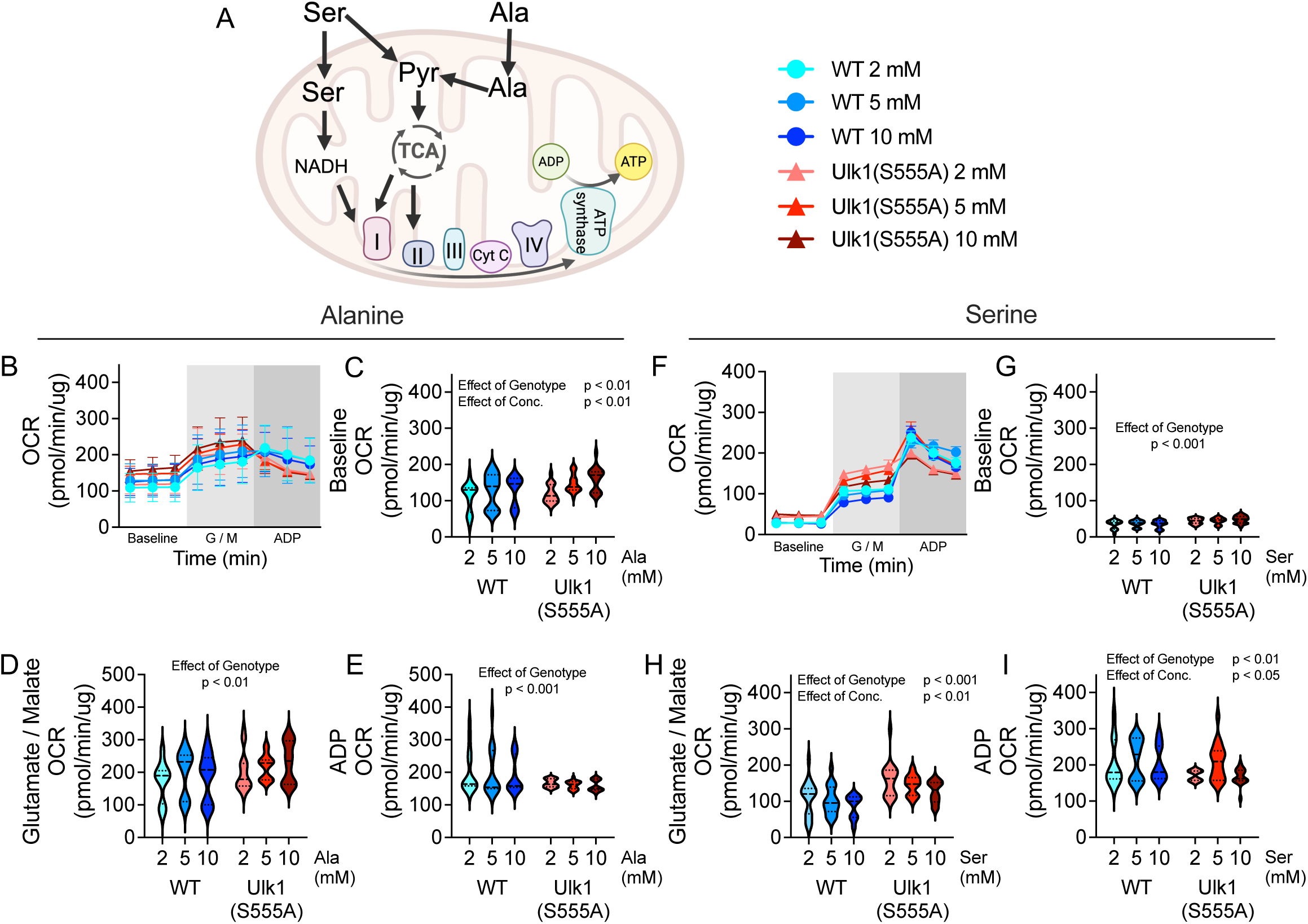
Loss of Ulk1(S555) increases mitochondrial oxidative capacity of alanine and serine in skeletal muscle. A) Illustration of the alanine and serine oxidation fates in mitochondria. B) Alanine incubated oxygen consumption rate (OCR) over time. C) Alanine incubated baseline OCR. D) Alanine incubated OCR with the addition of glutamate/malate, E) Alanine incubated OCR with the addition of 5 mM ADP. F) Serine incubated OCR over time. G) Serine incubated baseline OCR. H) Serine incubated OCR with the addition of glutamate/malate. I) Serine incubated OCR with the addition of 5 mM ADP. n=6 per group. Data presented as mean ± SEM and analyzed via two-way ANOVA.

### Evidence for increased amino acid transamination in liver with CR in Ulk1(S555A) mice disrupts mitochondrial respiration in the presence of pyruvate

Metabolic flexibility during nutrient stress occurs, in part, through coordination between skeletal muscle as the primary utilizer of glucose and liver as the primary site for gluconeogenesis via amino acids (Felig, 1973; Hewton et al., 2021; Sarabhai & Roden, 2019). Transamination of amino acids, such as alanine, has been demonstrated to be elevated when glucose metabolism is impaired, as occurs in Type II Diabetes (Okun et al., 2021). As glucose oxidation capacity following CR was also impaired in the liver of Ulk1(S555A) mice (Fig. 1J), we examined liver of Ulk1(S555A) mice following CR via untargeted metabolomics. Seven of the top ten upregulated pathways in liver via Metabolic pathway impact analysis were related to amino acid metabolism in Ulk1(S555A) mice following CR (Fig. 4A, B; S3B), consistent with increased transamination of amino acids. Analysis of hepatic amino acids showed elevated levels of glucogenic and ketogenic amino acids that were either elevated in Ulk1(S555A) and unaffected by CR (e.g. glycine, methionine, and lysine, with serine nearing significance), further elevated in response to CR (e.g. aspartic acid, glutamate, glutamine, methionine, and tyrosine), or inversely affected by CR compared to WT (e.g. alanine, threonine, and taurine) (Fig. S3C). Given the impaired oxidative capacity of glucose in Ulk1(S555A) mice following CR (Fig. 1 H-J), it’s possible that the liver becomes biased towards amino acid transamination to meet energetic demand.

**Figure 4.**
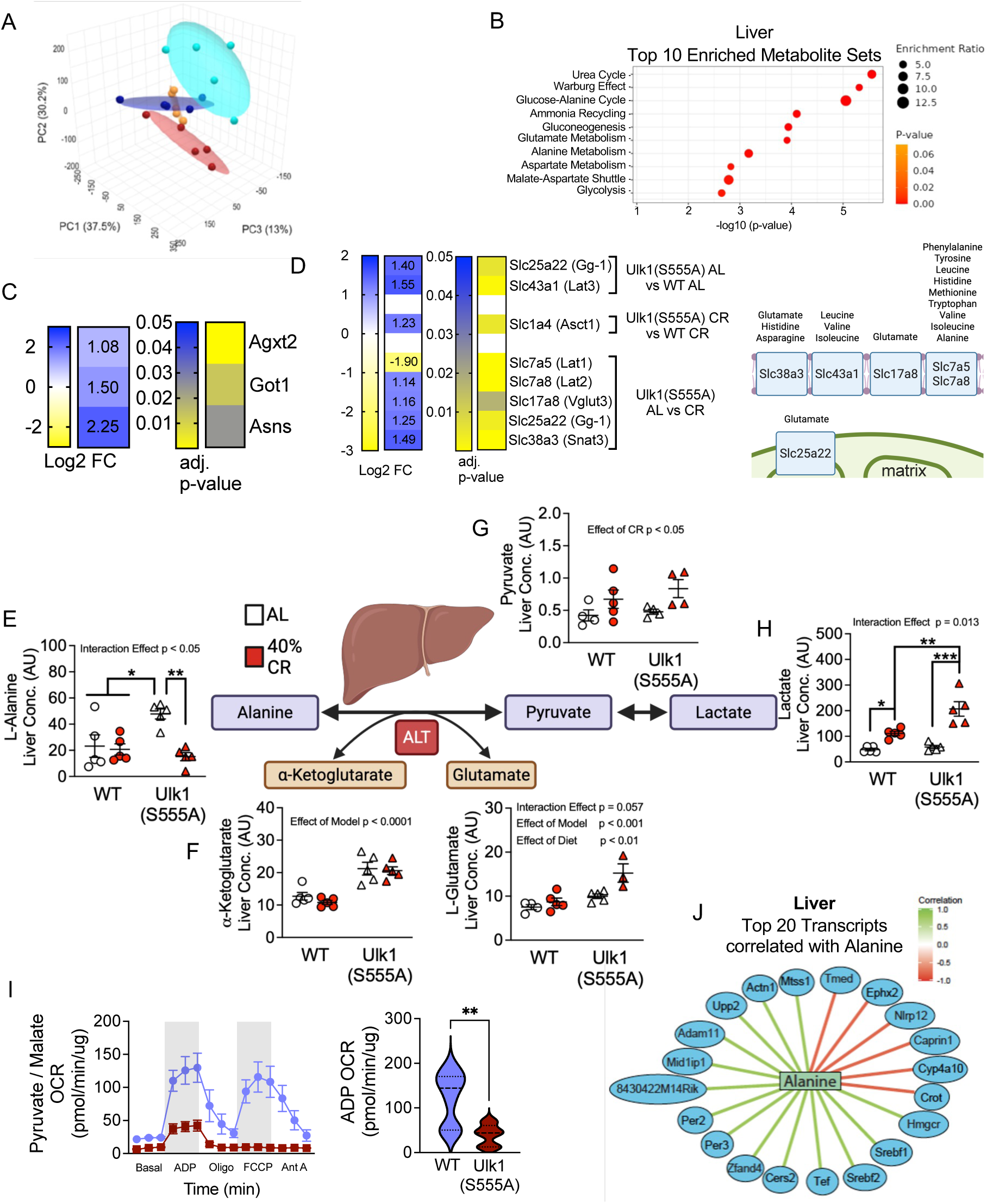
Caloric restriction leads to evidence of increased amino acid transamination in liver and inhibition of mitochondrial respiration in the presence of pyruvate. A) PCA plot of liver metabolites. B) Enriched metabolites in liver of Ulk1(S555A) with CR. C) Significantly upregulated genes in the Alanine, Aspartate, and Glutamate Metabolism pathway in liver of AL Ulk1(S555A) mice. D) Significantly unregulated genes for amino acid transporters in liver. E-H) Alanine transamination pathway and metabolite analysis. I) Pyruvate incubated liver mitochondria OCR. J) Mixomics analysis of liver transcripts correlated with alanine. N = 5 per group. Data presented as mean ± SEM and analyzed via two-way ANOVA with post-hoc analysis when a significant interaction effect was found (* = p < 0.05, ** = p < 0.01).

To gain insight into whether liver from Ulk1(S555A) mice may have increased capacity to transaminate amino acids, we performed bulk RNA sequencing (RNAseq). Ulk1(S555A) AL mice positively correlated with Alanine, Aspartate, and Glutamate Metabolism (Fig. S4A), which coincided with increased expression of genes involved in amino acid transamination (Fig. 4C). Agxt2 encodes a mitochondrial enzyme that produces glycine. Got1 encodes glutamate oxaloacetate transaminase 1, which catalyzes the conversion of α-ketoglutarate to glutamate in the cytosol. Asns encodes for asparagine synthetase, creating asparagine and glutamate from aspartic acid. This evidence for increased capacity for amino acid transamination in the liver of Ulk1(S555A) mice was further supported by increased expression of amino acid transport genes (Fig. 4D). In sum, impaired capacity to oxidize glucose in the absence of Ulk1(S555) may result in increased transamination of amino acids in the liver.

As alanine is well understood to be transaminated in the liver during prolonged fasting (Sarabhai & Roden, 2019), and elevated liver alanine in Ulk1(S555A) mice was significantly decreased following CR (Fig. 4E), we focused on analyzing metabolites associated with alanine transamination. The CR-dependent decline of liver alanine in Ulk1(S555A) mice (Fig. 4E) was accompanied by maintained α-ketoglutarate and elevated glutamate (Fig. 4D), supportive of increased transamination of alanine. Subsequently, pyruvate was elevated in both WT and Ulk1(S555A) liver with CR (Fig. 4F), but Ulk1(S555A) mice had significantly more lactate compared to WT (Fig. 4G, H). In alignment with the Warburg effect noted in the metabolic pathway analysis (Fig. 4B), elevated lactate in liver of Ulk1(S555A) mice following CR (Fig. 4H) suggests impaired conversion of pyruvate in liver mitochondria in the absence of Ulk1(S555). To test whether liver mitochondrial utilization of pyruvate was impaired in Ulk1(S555A) mice, hepatic mitochondrial respiration was assessed in isolated mitochondria incubated in pyruvate/malate following an overnight fast. Compared to WT, Ulk1(S555A) hepatic mitochondrial OCR remained significantly suppressed throughout the respiration assay (Fig. 4I, S4B, C). This may suggest increasing emphasis on amino acid transamination in the liver, presumably for export for use by other tissues such as skeletal muscle (Sarabhai & Roden, 2019), is at the expense of itself.

To gain insight into how increased transamination of amino acids in the liver in Ulk1(S555A) mice with CR reprograms liver metabolism, we performed mixOmics analysis between liver metabolomic and bulk RNAseq datasets, using alanine as our proxy metabolite. Liver alanine negatively correlated with fatty acid oxidation genes, *Cyp4a10* and *Crot* (Lin et al., 2006; Sanford et al., 2023), and positively correlated with cholesterol biosynthesis genes, *Srebf1/2* and *Hmgcr* (Daniels et al., 2023; Mteyrek et al., 2016; Wang et al., 2020) (Fig. 4J). Liver alanine was also positively correlated with circadian rhythm genes, *Per2/3* (Fig. 4J), consistent with impaired metabolic flexibility (Reinke & Asher, 2016). In sum, phosphorylation of Ulk1 at S555 is required for metabolic adaptation to CR, which may bias metabolism towards amino acids over glucose, suggesting Ulk1(S555) is an important metabolic switch point.

### Altered metabolic switch during fasting in Ulk1(S555A) mice

Mice on CR, where the daily allocation of food is given all at once, as we performed, eat all provided food within a few hours (Acosta-Rodriguez et al., 2017; Duregon et al., 2023; Mitchell et al., 2019). Therefore, self-imposing a daily, prolonged (20-22 hrs) fasted period. As Ulk1(S555A) mice did not increase their capacity to oxidize glucose within skeletal muscle and liver following CR (Fig. 1H-J) and liver MixOmics analysis suggested potential alterations in metabolic flexibility in relation to alanine (Fig. 4J), we hypothesized loss of Ulk1(S555) *in vivo* impaired metabolic switching during the daily self-imposed fasted window (Acosta-Rodriguez et al., 2017; Duregon et al., 2023; Mitchell et al., 2019). WT and Ulk1(S555A) mice were monitored in metabolic cages for 24 hours with *ad libitum* access to food, then fasted for 24 hours (Fig. 5A). There was no difference in respiratory exchange ratio (RER) during either light or dark phases when food access was unrestricted (Fig. 5A-B). Once food was removed, RER of WT mice dropped below 0.75 within 6 hours of fasting (Fig. 5A), indicative of increased reliance on fat as a substrate. However, after an initial drop upon fasting, which may have been due to opening cages to remove food, Ulk1(S555A) mice RER did not drop below 0.75 until after ∼18 hours of fasting (Fig.5A), indicating maintained reliance on mixed substrates. Along with the delayed metabolic switch, Ulk1(S555A) mice were less active, while energy expenditure and heat were equal between groups (Fig. 5B, Fig. S5A, B).

**Figure 5.**
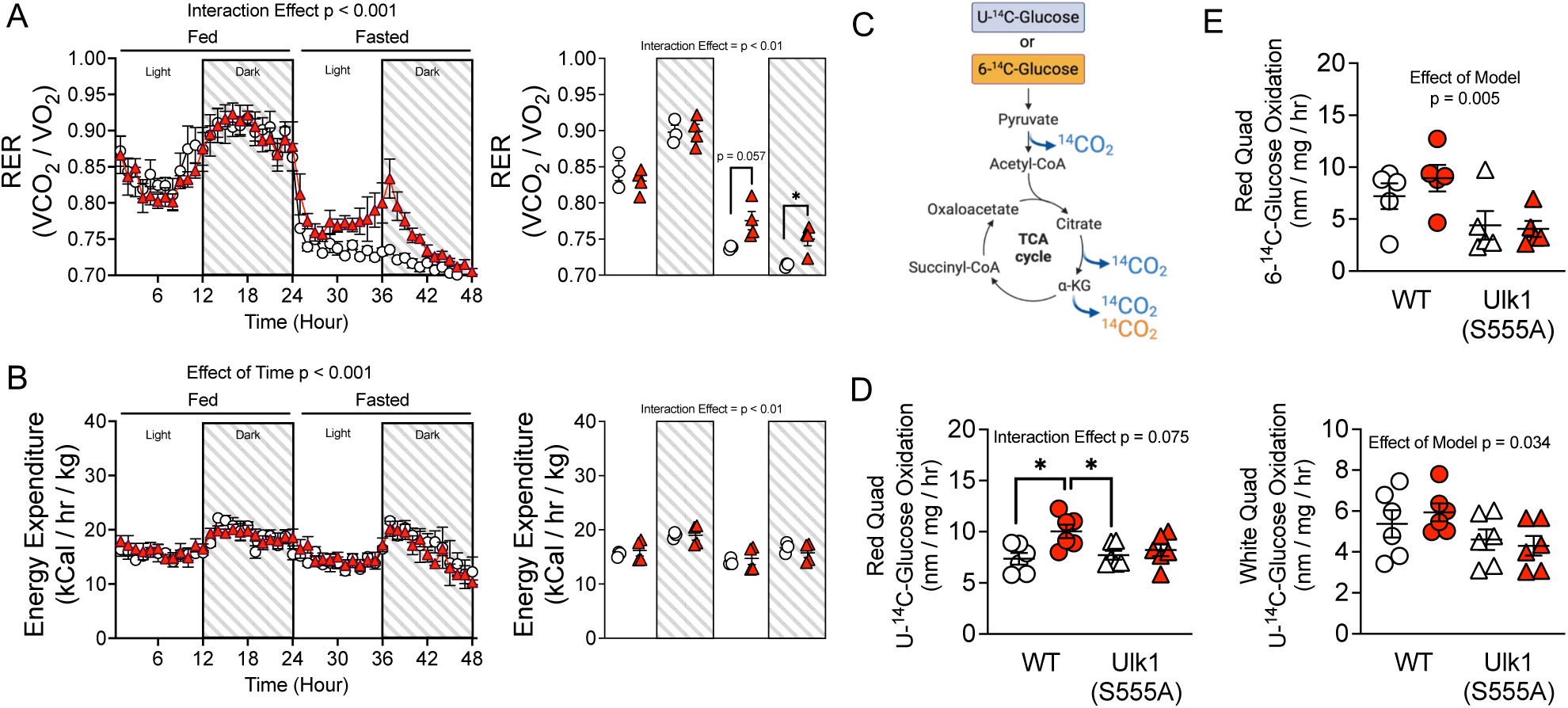
Altered metabolic switch during fasting in the absence of Ulk1(S555). A) WT and Ulk1(S555A) 48-hour respiratory exchange ratio (RER). B) WT and Ulk1(S555A) 48-hour energy expenditure (EE). C) U-^14^C-Glucose and 6-^14^C-Glucose oxidation pathway. D) U-^14^C-Glucose oxidation in red and white fibers isolated from quadricep skeletal muscle in fed and 24-hour fasted states. E) 6-^14^C-Glucose oxidation in red fibers isolated from quadricep skeletal muscle in fed and 24-hour fasted states. N = 4-5 per group. Data presented as mean ± SEM and analyzed via repeated measures or two-way ANOVA with post-hoc analysis when a significant interaction effect was found (* = p < 0.05).

To determine if glucose oxidation capacity was also impaired during acute fasting, we utilized U-^14^C-Glucose and 6-^14^C-Glucose to compare the fate of all carbons in glucose vs. those that are oxidized last in the TCA cycle, respectively (Fig. 5C). Capacity to oxidize glucose *ex vivo* via U-[^14^C]-Glucose (Fig. 5C) increased in red quadricep muscle fibers from WT in response to a 24-hour fast at 3 and 9 months of age but was unchanged by fasting in Ulk1(S555A) mice (Fig. 5D; Fig. S5C). A genotype effect was also observed in white quadricep muscle fibers following 24-hour fasting (Fig. 5D). Further, Ulk1(S555A) mice had lower capacity to oxidize 6-^14^C-Glucose in red quadricep muscle fibers from Ulk1(S555A) (Fig. 5E). Put together, the lack of increased oxidative capacity of U-^14^C-Glucose and 6-^14^C-Glucose supports a preferential shift away from glucose during a prolonged fast to maintain energetic homeostasis when Ulk1 is unable to be phosphorylated at S555. As RER was reflective of mixed substrate use until after ∼18 hours of fasting (Fig. 5A), these findings may collectively reflect a reliance on amino acids in Ulk1(S555A) mice (Figs. 2-4) and/or diversion of substrates to futile cycles. A reliance on amino acids could explain why Ulk1(S555A) mice were significantly less active despite energy expenditure being equivalent (Fig. 5B, Fig. S5A), as protein breakdown and urea synthesis are energetically costly (Hall, 2006; Veldhorst et al., 2009), leading to the elevation of basal energy expenditure with a lack of increased physical activity.

### Impaired Ulk1(S555) phosphorylation under nutrient stress and autophagy-derived amino acids

During fasting, skeletal muscle relies on the supply of nutrient substrates delivered through circulation (Sarabhai & Roden, 2019). To gain insight into preferred substrates in Ulk1(S555A) mice during fasting, we performed untargeted metabolomics of plasma following 24-hour fasting. In Ulk1(S555A) mice, circulating urea was elevated with fasting (Fig. 6A-B), coinciding with elevated circulating branched chain amino acids (BCAA) as well as glutamine (Fig. 6C-E). This finding suggests elevated amino acid metabolism and proteolysis, respectively (Flakoll et al., 1989; Paulusma et al., 2022; Remesy et al., 1997), which was further supported by metabolic pathway analysis of the fasted plasma metabolome in Ulk1(S555A) mice (Fig. S6) and evidence of muscle-dependent atrophy with fasting or CR (Fig. S7A-I).

**Figure 6.**
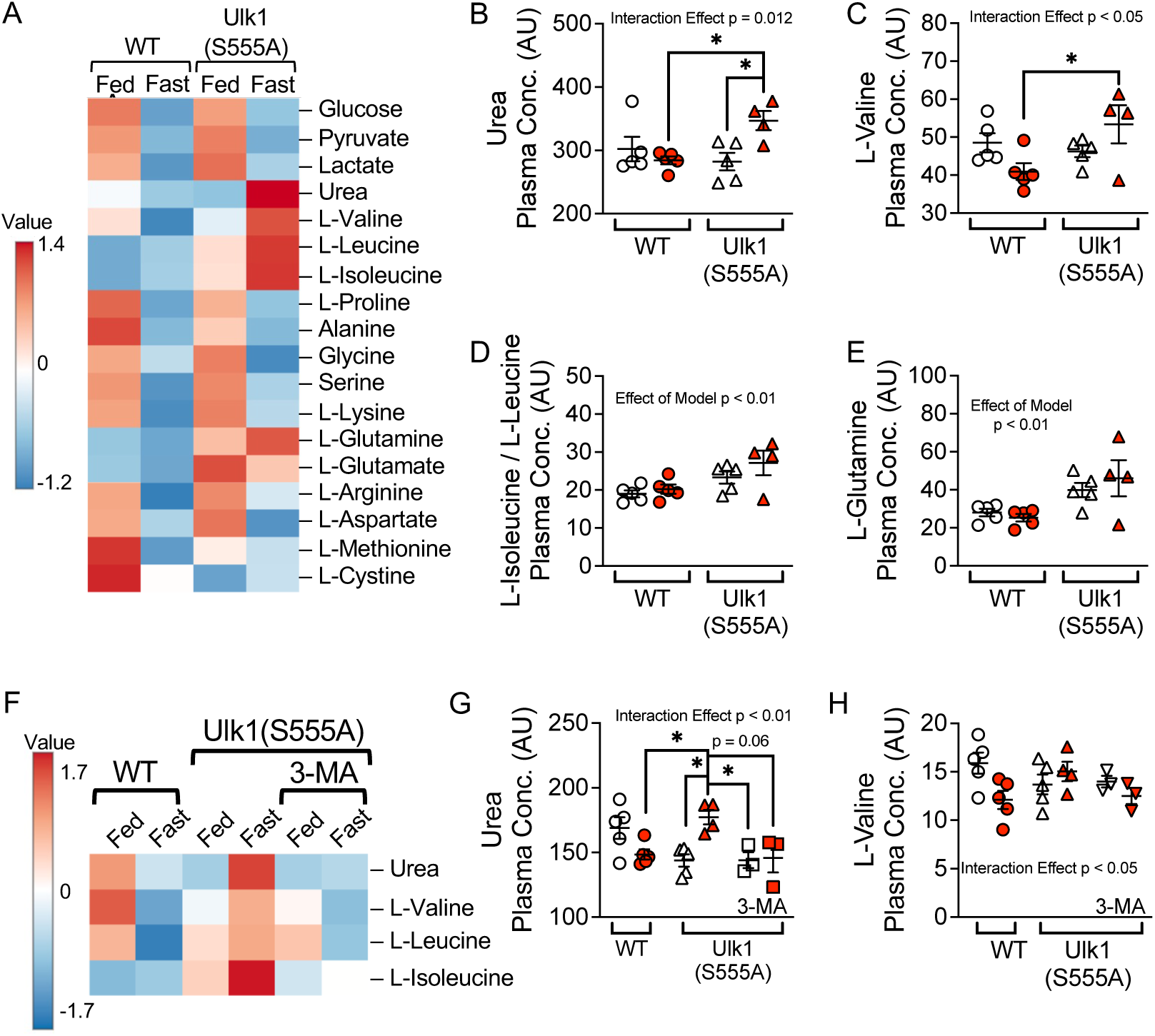
Increased plasma urea and branched chain amino acids with fasting in Ulk1(S555A) is attenuated with the autophagy inhibitor 3-MA. A) Plasma metabolomics heatmap of amino acids in WT and Ulk1(S555A) mice in fed and 24-hour fasted states. B) Plasma Urea, C) Valine, and D) Isoleucine/leucine. E) Plasma metabolomics heatmap of urea and BCAA following 3-MA administration in Ulk1(S555A) mice. F) Plasma Urea and G) Valine. n = 3 - 5 per group. Data presented as mean ± SEM and analyzed via two-way ANOVA with post-hoc analysis when a significant interaction effect was found (* = p < 0.05).

Recent evidence has challenged the paradigm that Ulk1(S555) phosphorylation initiates its autophagy functions (Kazyken et al., 2024; Park et al., 2023). That Ulk1(S555) would restrain autophagy is consistent with sustained LC3II levels in heart of Ulk1(S555A) mice (Guan, Zhang, et al., 2024) and elevated skeletal muscle mitophagy (Guan, Spaulding, et al., 2024). Therefore, we hypothesized sustained autophagy in Ulk1(S555A) mice sustains circulating area and BCAA levels. To test this, we treated Ulk1(S555A) mice with 3-methyladenine (3MA) via intraperitoneal injection once daily for three days before a 24-hour fast. 3MA disrupts the interaction between Ulk1 and Vsp34, inhibiting formation of the Ulk1 complex and autophagy (Galluzzi et al., 2017). Pre-treatment with 3MA attenuated the fasting-induced increase in circulating urea and BCAA in Ulk1(S555A) mice (Fig. 6E-G; S7E-G). Taken as a whole, these findings suggest phosphorylation of Ulk1 at S555 is required for metabolic switching and suggest impairment in Ulk1(S555) results in an increased reliance on autophagy-derived amino acids as energetic substrates to maintain energetic homeostasis (Fig. 6H).

## Discussion

Inter-organ communication is critical for maintaining metabolic flexibility—the ability to adapt substrate source to changes in substrate availability (Goodpaster & Sparks, 2017). In healthy states, the body can efficiently transition between fuel sources depending on demand. However, in metabolic disorders and aging, inter-organ communication breaks down, impairing fuel switching and promoting metabolic inflexibility (Castillo-Armengol et al., 2019; Goodpaster & Sparks, 2017). Skeletal muscle accounts for 60-80% of glucose utilization (Ng et al., 2012), accentuating its central role in regulating whole-body glucose homeostasis. As circulating glucose levels fall during fasting, skeletal muscle transiently shifts to amino acid catabolism, primarily utilizing amino acids as gluconeogenic energy sources. Amino acids are also released and taken up by the liver, converted to pyruvate or acetyl-CoA, and used in gluconeogenesis, thereby preserving plasma glucose levels for glucose-dependent tissues (Felig, 1973). As fasting progresses, both liver and muscle engage in coordinated metabolic reprogramming: the liver increases fatty acid oxidation and ketogenesis to produce alternative fuels, while skeletal muscle upregulates fatty acid uptake and β-oxidation, transitioning away from amino acid catabolism to conserve muscle protein. This adaptive inter-organ communication allows for efficient substrate switching—glucose to amino acids to lipids—ensuring that energy needs are met without depleting critical reserves. Our findings in the current study suggest phosphorylation of Ulk1 at S555 facilitates fuel switching during nutrient stress, highlighting a critical role in maintaining metabolic flexibility and potential target against metabolic dysfunction.

There are four known Ulk variants, with Ulk1 and Ulk2 being the most similar, with 55% homology (Chan et al., 2009; Hara et al., 2008). However, insulin-induced phosphorylation of Akt in skeletal muscle is impaired following exercise training in tamoxifen-inducible Ulk1 knockout mice, but not Ulk2 knockouts (Drake et al., 2021; Laker et al., 2017), suggesting Ulk1 is specifically needed for adaptation of glucose metabolism to energetic stress *in vivo*. In cell culture, Ulk1 has been shown to directly phosphorylate the glycolytic enzymes HK, thus promoting glucose uptake, and FBP1, continuing glycolysis over gluconeogenesis (Li et al., 2016). Further, Ulk1 has also been shown to phosphorylate PFK1 and ENO1, thus inhibiting their activity, causing the accumulation of glucose-6-phosphate, enhancing NADPH production, and maintaining redox homeostasis (Li et al., 2016). Our data extend these findings in vivo by demonstrating Ulk1 phosphorylation at S555 is the post-translational modification required for Ulk1-mediated regulation of glucose oxidation during nutrient stress in skeletal muscle and liver, possibly resulting in increased reliance on amino acids.

Under normal conditions, the reliance on amino acids as an energetic substrate is either brief, as during the transition from glucose to fat, or as a last resort following prolonged fasting (Petersen et al., 2019). However, our findings suggest that in Ulk1(S555A) mice, reliance on amino acids as an energetic substrate is prolonged (∼12 hours) during a 24-hour fast and a major energy source during short-term CR. Ulk1(S555) was recently demonstrated to be sensitive to amino acid availability in a variety of cell lines, with amino acid restriction increasing S555 phosphorylation and the addition of amino acids serine, hydroxyproline, and glutamine decreasing S555 phosphorylation (Kazyken et al.). In glucose-deprived HEK293 cells, protein synthesis is suppressed, and the essential amino acid leucine is made available as an energetic substrate through Ulk1-dependent phosphorylation of leucyl-tRNA synthetase 1 (LARS1) to maintain energetic homeostasis (Yoon et al., 2020). Ulk1(S555) may therefore act as an amino acid sensing mechanism that dictates the transition from glucose to fat.

We recently showed exercise-induced mitophagy is intact in Ulk1(S555A) mice (Guan, Spaulding, et al., 2024), which echoes other recent work suggesting Ulk1(S555) does not induce autophagy (Kazyken et al., 2024; Park et al., 2023). These recent studies are contrary to the previous model for Ulk1(S555) (Bujak et al., 2015; D. Egan et al., 2011; Laker et al., 2017). Here, we present evidence that elevated amino acid metabolism in Ulk1(S555A) mice, as evidenced by increased plasma urea, is dependent on sustained autophagy; reinforcing the notion that phosphorylation of Ulk1 at S555 inhibits autophagy. Taken together, the inability of Ulk1(S555A) mice to enhance glucose oxidation in skeletal muscle and liver following different periods of nutrient stress suggests Ulk1 phosphorylation at S555 controls metabolic flexibility, in part, through modulating autophagy.

Amino acids and glucose can suppress the utilization of each other to promote either anabolic or catabolic processes (Shao et al., 2018). Thus, reliance on amino acids as a substrate during nutrient stress in Ulk1(S555A) mice may also contribute to impaired capacity to oxidize glucose. For example, pharmacologically inhibiting mitochondrial complex I-mediated respiration elevates serine catabolism, leading to an accumulation of NADH, suppressing other metabolic pathways known to produce NADH (glucose, glutamine, and fat oxidation) (Yang et al., 2020). Alternatively, serine-derived NADH may be an essential adaptive response to maintain mitochondrial energetics (Lucas et al., 2018). Alanine is predominantly released and synthesized in skeletal muscle (Felig, 1973), while the liver is the primary site for alanine transamination to pyruvate (Hewton et al., 2021). Notably, our metabolomics data suggest Ulk1(S555A) AL mice exhibit elevated alanine concentrations in both skeletal muscle and liver, coupled with increased hepatic alanine transamination in liver with CR. In skeletal muscle mitochondria, Ulk1(S555A) mice had an affinity to oxidize alanine and serine to support basal and complex I-mediated respiration, highlighting a shift in mitochondrial substrate preference that may reflect adaptive mechanisms when Ulk1 cannot be phosphorylated at S555. Alanine has been shown to outcompete glucose as a substrate to fuel mitochondrial respiration in pancreatic ductal adenocarcinoma (PDAC) (Sousa et al., 2016). While loss of Ulk1(S555) may directly impair glycolysis through direct interruption of glycolysis (Li et al., 2016), preference for amino acids over glucose may further disrupt metabolic switching by outcompeting glucose-derived pyruvate and/or NADH to fuel mitochondrial metabolism. However, the inefficiency of amino acid oxidation may limit mitochondrial respiration during robust or prolonged nutrient stress (Hall, 2006; Veldhorst et al., 2009).

In the present study, we showed Ulk1(S555A) mice have impaired metabolic flexibility, reduced glucose oxidative capacity, and evidence of increased reliance on autophagy-derived amino acids. This altered substrate preference reflects metabolic features of insulin resistance (Krebs et al., 2002; Wang et al., 2011) and may contribute to the development of obesity-related disorders and type 2 diabetes. Furthermore, increased amino acid catabolism in the context of compromised glucose oxidation may predispose to sarcopenia and muscle loss, particularly since Ulk1(S555) may decline with age (Nichenko et al., 2021). Future studies should investigate disease-dependent relevance of Ulk1(S555) as a therapeutic target to promote metabolic flexibility.

## Materials and Methods

### Animals

All experiments and procedures were approved by the Institutional Animal Care and Use Committee (IACUC) at Virginia Tech. *Ulk1(S555A)* knock-in mice were generated in C57BL/6J background were generated at the Genetically Engineered Murine Model (GEMM) core facility at the University of Virginia (Guan, Spaulding, et al., 2024). Homozygotic knock-in mice were generated from homozygous breeding pairs in-house and compared to background controls (JAX). Mice were housed in temperature-controlled (21°C) quarters with a 12:12 h light-dark cycle and *ad libitum* access to water and chow (Purina) until designated for study.

### Caloric Restriction

At 6 months of age (24 weeks of age), mice were designated to *ad libitum* fed or caloric restriction (CR). Mice designated for CR were maintained in cages of 4-5 mice to normalize competition for food between groups. To calculate daily food allotment, food consumption was monitored for three consecutive days and averaged. Pre-CR NMR and GTT were taken (described below). For CR, mice were given food once daily at 1600. At the onset, CR groups spent 1 week at 20% CR as a step down and then continued at 40% CR for the subsequent 8 weeks. Post-CR NMR and GTT were taken during weeks 6 and 7, respectively, of CR to avoid any confounding with tracer or metabolomic measures.

### 24-hour Fast

Mice were designated into fed or 24-hour fast groups. Fed groups had ad libitum access to food until sacrificed. 24-hour fast groups were placed into new cages without food access for 24 hours before sacrifice. All harvests began at the onset of the light cycle (7:00).

### 3-Methlyadenine administration

Ulk1(S555A) mice were designated into fed or 24-hour fast groups. 3MA was administered via IP injection at a 30 mg/kg concentration for 3 consecutive days. Fed groups had ad libitum access to food until sacrificed. 24-hour fast groups were placed into new cages without food access for 24 hours before sacrifice. All harvest began at the onset of the light cycle (0700).

### Glucose tolerance

Glucose tolerance was performed at approximately 0800. For AL mice, food was removed at approximately 1800 to normalize the fasting period across groups. Blood glucose from the tail vein was measured before and following intraperitoneal injection of sterilized d-glucose (2mg/g) at 30, 60, and 120 min as previously (Drake et al., 2021; Guan, Spaulding, et al., 2024; Laker et al., 2017).

### Nuclear Magnetic Resonance

NMR (Bruker Minispec) was utilized to assess body composition. Before the assessment, the mini spec performed a quality control check of internal voltages, temperature, magnets, and NMR parameters using the manufacturer’s standards. Mice were placed and immobilized in a plastic tube and placed into the sample chamber for ∼2 minutes, or the duration of the scan.

### Calorimetry

Mice were individually housed in Columbus Comprehensive Lab Animal Monitoring System (CLAMS) metabolic cages and acclimated before a 48-hour observation (24-hour ad libitum fed/24-hour fasted) to assess respiratory exchange ratio, energy expenditure, and activity.

### Tissue Procurement

For the CR study, prior to tissue harvest, CR mice were fed at 1600, and then all mice had food removed at 1800 by placing them in clean cages without food. Beginning at approximately 0700, mice were anesthetized with 3% isofluorane. The quadriceps and liver were collected antemortem and divided, with one part for metabolomics, another part prepared for radio-labeled tracing, and the rest frozen in LN_2_ for later analyses. The metabolomics-designated tissues were clamp-frozen immediately and placed in LN_2_. The tracing-designated tissues were placed in phosphate-buffered saline (PBS) and immediately prepped for radio-labeled tracing. Additional tissues collected were done post-mortem.

### *Ex vivo* Radio-labeled glucose tracing

The liver was surgically dissected (above), and the quadriceps were further dissected into red (oxidative) and white (glycolytic) regions. Approximately 15-50mg of tissue was placed in 200 µl of modified sucrose EDTA medium (SET buffer) containing 250 mM sucrose, 1 mM EDTA, 10 mM Tris-HCl, and 1 mM ATP at pH 7.4. Tissues were minced with scissors, then SET buffer was added to create an approximate 20-fold dilution (weight : volume). Samples were homogenized in a Potter-Elvehjem glass homogenizer for 10 passes over 30 s at 150 rpm with a Teflon pestle. and then incubated in U-^14^C-glucose or 6-^14^C-glucose (American Radiolabeled Chemicals, St. Louis, MO, USA). Production of ^14^CO_2_ was assessed by scintillation counting (4500 Beckman Coulter) and interpreted for glucose oxidative capacity. Total protein content in tissue homogenates was measured via BCA (Thermo Fisher Scientific, Waltham, MA, USA) and used to normalize oxidation values.

### Untargeted Metabolomics (GC-MS)

Untargeted GC/MS-based metabolomics analysis was performed on skeletal muscle (quadriceps) and liver from Ulk1 (S555A) and WT mice. Metabolites were extracted with 90% methanol, dried, and derivatized. Metabolomic profiling was performed in an electron-ion (EI) mode using a Shimadzu triple quadrupole GCMS-TQ8050 NX (GC-MS/MS). Data analysis relied upon a combination of MS-Dial and MetaboAnalyst to identify features that were most different between groups (PCA plot, volcano plot, and heat map).

### RNA seq/Transcriptomics

Frozen liver and quadricep muscle were homogenized in LN_2_ via mortar and pestle, then added to Trizol. Total RNA was isolated via Qiagen RNeasy Mini Kit (74106). For RNA sequencing (Novogene, Durham, NC, USA), mRNA was purified from total RNA using poly T oligo-attached magnetic beads. Following fragmentation, first-string cDNA was synthesized using random hexamer primers, and second-strand cDNA was synthesized using dTTP for a non-directional library and checked with Qubit and real-time PCR for quantification and a bioanalyzer for size distribution detection. Libraries were pooled and sequenced on the Illumina platform (NovaSeq X plus series) according to library concentration and data amount (9 G per sample). Reference genome was built using Hisat2 v2.0.05 and differential expression analysis was performed using DESeq2Rpackage (1.20.0).

### MixOmics

To identify correlations between metabolomics and transcriptomics data, regularized Canonical Correlation Analysis (rCCA) was used (González et al., 2009). rCCA identifies pairs of linear combinations, canonical variates, that maximize the correlation between the two datasets under regularization constraints to prevent overfitting. Data was input as normalized fragments per kilobase of transcript per million mapped reads (FPKM) for transcriptomics data, while peak quantification data from MetaboAnalyst was used as-is for metabolomics data (see Untargeted Metabolomics (GC-MS) section, above). Transcripts with more than two zero or missing values in any single sample group were excluded. The rcc() function from the MixOmics R package was used to generate correlation structures, which were evaluated based on the strength of canonical correlations and the interpretability of component loadings (Rohart et al., 2017). Specific metabolites of interest were examined to identify the most strongly correlated transcripts, visualized via network plots.

### Mitochondrial isolation

Quadricep skeletal muscle and liver tissues were dissected and immediately placed in ice-cold Isolation Buffer Mitochondria-1 (IBM-1). Tissues were minced for 2 minutes, incubated with 0.25% trypsin in IBM-1 for 30 minutes on ice, and then centrifuged at 200 x g for 3 minutes at 4°C. The pellet was resuspended in IBM-1 and homogenized in a pre-chilled glass homogenizer (10 passes at 80 rpm). Homogenates were cleared by centrifugation at 700 x g for 10 minutes at 4°C. The resulting supernatant was filtered through a 100μm nylon mesh and transferred to chilled high-strength glass centrifuge tubes. Mitochondria were pelleted at 8,000 x g for 10 minutes at 4°C, resuspended in IBM-2, and re-pelleted under the same conditions. Final mitochondrial pellets were gently resuspended in 100μL IBM-2 and kept on ice. Protein concentrations were quantified using a BCA assay with 1:10 and 1:20 sample dilutions.

### Mitochondrial respiration

Mitochondrial respiration was measured using the Seahorse XFe96 extracellular flux analyzer (Agilent). Liver and skeletal muscle mitochondrial fractions were isolated from n = 6 WT and Ulk1(S555A) mice across three separate days. The six biological replicates were combined in pairs to create n = 3 biological duplicates and analyzed as six (Fig. 3) or three (Fig. 4I) technical replicates per pairing and condition. Mitochondrial-enriched lysates were seeded into the cell plate at a 2 ug/ul concentration. Media was prepared at the following concentrations: pyruvate (10 mM)/malate (5 mM) 7.4pH, succinate (10 mM) / rotenone (2 uM) 7.4pH, alanine (2 mM, 5 mM, 10 mM)/malate (5 mM) 7.4pH, and serine (2mM, 5mM, 10mM)/malate (5mM) 7.4pH. Ports were prepared with A: glutamate (10 mM) / malate (5 mM) and B: ADP (4 uM). Oxygen consumption rate (OCR) was obtained and calculated as previously described (10, 14). Mitochondrial respiration measurement cycles consisted of 3 min mixing, 2 min wait, and 3 min measurement cycles.

### Western Blotting

Total protein lysates were subjected to sodium dodecyl sulfate-polyacrylamide gel electrophoresis and transferred onto a nitrocellulose membrane. Briefly, tissues were electrically homogenized in ∼1 mL fractionation (FRAC) buffer [20 mM HEPES, 250 mM Sucrose, 0.1 mM EDTA, plus protease (Roche Diagnostics) and phosphatase (Sigma) inhibitors]. Whole-tissue lysate was resuspended in 50uL of Laemmli sample buffer, boiled for 5 min at 97°C, then frozen at −80°C until further analysis. Protein concentration was determined via RCDC assay. Membranes were probed with the following primary antibodies at 1:1000 dilution, targeting pUlk1 (S555) (CST #5869), tUlk1 (Sigma #A-7481), α-tubulin (Abcam #ab7291)

### Statistics

Data are presented as the mean ± SEM and were analyzed and plotted via GraphPad Prism v10. Data were analyzed via Student’s t-test when one variable was present, two-way ANOVA when two variables were present, or repeated measures two-way ANOVA when two variables were measured in the same animals over time. Post-hoc analyses were only performed when a significant interaction between a categorical and a quantitative variable was found, which is indicated in the figures where applicable. Statistical significance was established a priori as P < 0.05.

## Funding

This work was supported by start-up funds from the College of Agriculture and Life Sciences and Department of Human Nutrition, Foods, and Exercise, as well as NIH grants (R01AG080731 and R00AG057825) from the National Institute on Aging (NIA) to JCD. NIH-NIAMS grant (R56AR083948) was used to support SMC. NIH-NIA grant (R00AG070104) was used to support CPN. TWR acknowledges support for this research provided by the Office of the Vice Chancellor for Research at the University of Wisconsin-Madison with funding from the Wisconsin Alumni Research Foundation. JSW was supported by the Seale Innovation Fund and NIH-NHLBI grant (R01HL156667).

## Acknowledgements

We thank Dr. Zhen Yan at the Center for Exercise Medicine Research at the Fralin Biomedical Research Institute at Virginia Tech for the generous gift of the Ulk1(S555A) mice. We also thank members of JCD’s laboratory for critical feedback and discussion.

## Author Contributions

Study Design: OSW, ASN, JCD. Conducting experiments: OSW, ASN, MHB, NA, SNH, DSB, JRB, BJK, KSJ, AKA, AVZ, STB, RPM, HZ, JCD. Analyzing data: OSW, ASN, SAT, CPN, SEC, TWR, JSW, JCD. Manuscript drafting: all authors.

## Conflict of Interest Statement

The authors have no conflicts to disclose.

## Data Availability

The data within this article will be made available following peer-reviewed publication.

## Supplemental Figure Legends

**Figure S1** A) Average food intake (g) over 3 days. B) NMR lean mass delta (%BM). C) NMR fat mass delta (%BM). D) Liver phospho-Ulk1(S555), total Ulk1, and ⍺-tubulin. E) Immunoblot quantification of liver pUlk1(S555)/total Ulk1. F) Immunoblot quantification of liver total Ulk1 / αTubulin. n = 5 per group. Data presented as mean ± SEM and analyzed via student’s t-test (A) or two-way ANOVA with post-hoc analysis when a significant interaction effect was found (** = p < 0.01, *** = p < 0.001).

**Figure S2** Skeletal muscle metabolomics. A) PCA plot generated in Metaboanalyst. B) Pathway impact of metabolites. C) Detected amino acid concentrations (glucogenic, ketogenic, both). D) Skeletal muscle citric acid and transcripts associated with glycolysis from bulk RNAseq. n=5. Data presented as mean ± SEM and analyzed via two-way ANOVA with post-hoc analysis when a significant interaction effect was found (** = p < 0.01).

**Figure S3** Liver metabolomics. A) PCA plot generated in Metaboanalyst. B) Pathway impact of metabolites. C) Detected amino acids concentrations in liver (glucogenic, ketogenic, both). n=5. Data presented as mean ± SEM and analyzed via two-way ANOVA with post-hoc analysis when a significant interaction effect was found (* = p < 0.05, ** = p < 0.01, *** = p < 0.001).

**Figure S4** A) Enrichment Scores of WT and Ulk1(S555A), AL and CR groups from Liver bulk RNAseq. B) WT and Ulk1(S555A) pyruvate incubated liver mitochondria OCR at baseline. C) WT and Ulk1(S555A) pyruvate incubated liver mitochondria OCR in presence of FCCP.

**Figure S5** A) Activity (m) during fed and 24-hour fasted periods. B) Heat (kcal/hour) during fed and 24-hour fasted periods. C) U-C-glucose oxidation pathway and oxidation in red fibers isolated from the quadriceps of 9-month-old animals. n = 3 - 4. Data presented as mean ± SEM and analyzed via two-way ANOVA with post-hoc analysis when a significant interaction effect was found (** = p < 0.01).

**Figure S6** WT and Ulk1(S555A) mice in *ad libitum* fed or 24-hour fasted. A) Plasma metabolomics enriched metabolite. n = 4 - 5. Data presented as mean ± SEM and analyzed via two-way ANOVA with post-hoc analysis when a significant interaction effect was found (* = p < 0.05, ** = p < 0.01).

**Figure S7** A-D) Tissue weights normalized to either tibia length or body mass from fed or 24 hour fasted WT and Ulk1(S555A) mice. E-I) Tissue weights normalized to either tibia length or body mass from AL or CR WT and Ulk1(S555A) mice. n=5. Data presented as mean ± SEM and analyzed via two-way ANOVA with post-hoc analysis when a significant interaction effect was found (* = p < 0.05, ** = p < 0.01, *** = p < 0.001).

**Figure S8** WT mice in an *ad libitum* fed or 24-hour fasted state and Ulk1(S555A) and Ulk1(S555A) 3MA mice in an *ad libitum* or 24-hour fasted state. A) Heatmap of Urea and branched-chain amino acids from plasma metabolomics. B) Plasma Leucine. C) Plasma Isoleucine. n = 3 - 5 per group. Data presented as mean ± SEM and analyzed via two-way ANOVA with post-hoc analysis when a significant interaction effect was found (* = p < 0.05).

## Notes

### Competing Interest Statement

The authors have declared no competing interest.

### Summary of Updates

addition of RNAseq analyses, liver mitochondria respiration, and plasma Metabolomics following treatment with the autophagy inhibitor 3-MA

## References

Acosta-Rodriguez, V. A., de Groot, M. H. M., Rijo-Ferreira, F., Green, C. B., & Takahashi, J. S. (2017). Mice under Caloric Restriction Self-Impose a Temporal Restriction of Food Intake as Revealed by an Automated Feeder System. Cell Metab, 26(1), 267–277 e262. 10.1016/j.cmet.2017.06.007

Bujak, A. L., Crane, J. D., Lally, J. S., Ford, R. J., Kang, S. J., Rebalka, I. A., Green, A. E., Kemp, B. E., Hawke, T. J., Schertzer, J. D., & Steinberg, G. R. (2015). AMPK activation of muscle autophagy prevents fasting-induced hypoglycemia and myopathy during aging. Cell Metab, 21(6), 883–890. 10.1016/j.cmet.2015.05.016

Castillo-Armengol, J., Fajas, L., & Lopez-Mejia, I. C. (2019). Inter-organ communication: a gatekeeper for metabolic health. EMBO Rep, 20(9), e47903. 10.15252/embr.201947903

Chan, E. Y., Longatti, A., McKnight, N. C., & Tooze, S. A. (2009). Kinase-inactivated ULK proteins inhibit autophagy via their conserved C-terminal domains using an Atg13-independent mechanism. Mol Cell Biol, 29(1), 157–171. 10.1128/MCB.01082-08

Chen, C. N., Lin, S. Y., Liao, Y. H., Li, Z. J., & Wong, A. M. (2015). Late-onset caloric restriction alters skeletal muscle metabolism by modulating pyruvate metabolism. Am J Physiol Endocrinol Metab, 308(11), E942–949. 10.1152/ajpendo.00508.2014

Daniels, L. J., Kay, D., Marjot, T., Hodson, L., & Ray, D. W. (2023). Circadian regulation of liver metabolism: experimental approaches in human, rodent, and cellular models. Am J Physiol Cell Physiol, 325(5), C1158–C1177. 10.1152/ajpcell.00551.2022

Drake, J. C., Wilson, R. J., Cui, D., Guan, Y., Kundu, M., Zhang, M., & Yan, Z. (2021). Ulk1, Not Ulk2, Is Required for Exercise Training-Induced Improvement of Insulin Response in Skeletal Muscle. Front Physiol, 12, 732308. 10.3389/fphys.2021.732308

Duregon, E., Fernandez, M. E., Martinez Romero, J., Di Germanio, C., Cabassa, M., Voloshchuk, R., Ehrlich-Mora, M. R., Moats, J. M., Wong, S., Bosompra, O., Rudderow, A., Morrell, C. H., Camandola, S., Price, N. L., Aon, M. A., Bernier, M., & de Cabo, R. (2023). Prolonged fasting times reap greater geroprotective effects when combined with caloric restriction in adult female mice. Cell Metab, 35(7), 1179–1194 e1175. 10.1016/j.cmet.2023.05.003

Duregon, E., Pomatto-Watson, L., Bernier, M., Price, N. L., & de Cabo, R. (2021). Intermittent fasting: from calories to time restriction. Geroscience, 43(3), 1083–1092. 10.1007/s11357-021-00335-z

Egan, D., Shackelfor, D., Mihaylova, M., Gelino, S., Kohnz, R., Mair, W., Vasquez, D., Joshi, A., Gwinn, D., Taylor, R., Asara, J., Fitzpatrick, J., Dillin, A., Viollet, B., Kundu, M., Hansen, M., & Shaw, R. (2011). Phosphorylation of ULK1 (hATG1) by AMP-activated protein kinase connects energy sensing to mitophagy. Science, 331, 456–461.

Egan, D. F., Shackelford, D. B., Mihaylova, M. M., Gelino, S., Kohnz, R. A., Mair, W., Vasquez, D. S., Joshi, A., Gwinn, D. M., Taylor, R., Asara, J. M., Fitzpatrick, J., Dillin, A., Viollet, B., Kundu, M., Hansen, M., & Shaw, R. J. (2011). Phosphorylation of ULK1 (hATG1) by AMP-activated protein kinase connects energy sensing to mitophagy. Science, 331(6016), 456–461. 10.1126/science.1196371

Felig, P. (1973). The glucose-alanine cycle. Metabolism, 22(2), 179–207. 10.1016/0026-0495(73)90269-2

Flakoll, P. J., Kulaylat, M., Frexes-Steed, M., Hourani, H., Brown, L. L., Hill, J. O., & Abumrad, N. N. (1989). Amino acids augment insulin’s suppression of whole body proteolysis. Am J Physiol, 257(6 Pt 1), E839-847. 10.1152/ajpendo.1989.257.6.E839

Fok, W. C., Zhang, Y., Salmon, A. B., Bhattacharya, A., Gunda, R., Jones, D., Ward, W., Fisher, K., Richardson, A., & Perez, V. I. (2013). Short-term treatment with rapamycin and dietary restriction have overlapping and distinctive effects in young mice. J Gerontol A Biol Sci Med Sci, 68(2), 108–116. 10.1093/gerona/gls127

Galluzzi, L., Bravo-San Pedro, J. M., Levine, B., Green, D. R., & Kroemer, G. (2017). Pharmacological modulation of autophagy: therapeutic potential and persisting obstacles. Nat Rev Drug Discov, 16(7), 487–511. 10.1038/nrd.2017.22

González, I., Déjean, S., Martin, P. G. P., Gonçalves, O., Besse, P., & Baccini, A. (2009). Highlighting relationships between heterogeneous biological data through graphical displays based on regularized canonical correlation analysis. Journal of Biological Systems, 17(02), 173–199. 10.1142/s0218339009002831

Goodpaster, B. H., & Sparks, L. M. (2017). Metabolic Flexibility in Health and Disease. Cell Metab, 25(5), 1027–1036. 10.1016/j.cmet.2017.04.015

Green, C. L., Lamming, D. W., & Fontana, L. (2022). Molecular mechanisms of dietary restriction promoting health and longevity. Nat Rev Mol Cell Biol, 23(1), 56–73. 10.1038/s41580-021-00411-4

Guan, Y., Spaulding, H., Yu, Q., Zhang, M., Willoughby, O., Drake, J. C., & Yan, Z. (2024). Ulk1 phosphorylation at S555 is not required for endurance training-induced improvements in exercise and metabolic capacity in mice. J Appl Physiol (1985), 137(2), 223–232. 10.1152/japplphysiol.00742.2023

Guan, Y., Zhang, M., Lacy, C., Shah, S., Epstein, F. H., & Yan, Z. (2024). Endurance Exercise Training Mitigates Diastolic Dysfunction in Diabetic Mice Independent of Phosphorylation of Ulk1 at S555. Int J Mol Sci, 25(1). 10.3390/ijms25010633

Hall, K. D. (2006). Computational model of in vivo human energy metabolism during semistarvation and refeeding. Am J Physiol Endocrinol Metab, 291(1), E23–37. 10.1152/ajpendo.00523.2005

Hara, T., Takamura, A., Kishi, C., Iemura, S., Natsume, T., Guan, J. L., & Mizushima, N. (2008). FIP200, a ULK-interacting protein, is required for autophagosome formation in mammalian cells. J Cell Biol, 181(3), 497–510. 10.1083/jcb.200712064

Hewton, K. G., Johal, A. S., & Parker, S. J. (2021). Transporters at the Interface between Cytosolic and Mitochondrial Amino Acid Metabolism. Metabolites, 11(2). 10.3390/metabo11020112

Kazyken, D., Dame, S. G., Wang, C., Wadley, M., & Fingar, D. C. (2024). Unexpected roles for AMPK in the suppression of autophagy and the reactivation of MTORC1 signaling during prolonged amino acid deprivation. Autophagy, 20(9), 2017–2040. 10.1080/15548627.2024.2355074

Krebs, M., Krssak, M., Bernroider, E., Anderwald, C., Brehm, A., Meyerspeer, M., Nowotny, P., Roth, E., Waldhausl, W., & Roden, M. (2002). Mechanism of amino acid-induced skeletal muscle insulin resistance in humans. Diabetes, 51(3), 599–605. 10.2337/diabetes.51.3.599

Laker, R. C., Drake, J. C., Wilson, R. J., Lira, V. A., Lewellen, B. M., Ryall, K. A., Fisher, C. C., Zhang, M., Saucerman, J. J., Goodyear, L. J., Kundu, M., & Yan, Z. (2017). Ampk phosphorylation of Ulk1 is required for targeting of mitochondria to lysosomes in exercise-induced mitophagy. Nat Commun, 8(1), 548. 10.1038/s41467-017-00520-9

Lee, M. B., Hill, C. M., Bitto, A., & Kaeberlein, M. (2021). Antiaging diets: Separating fact from fiction. Science, 374(6570), eabe7365. 10.1126/science.abe7365

Li, T. Y., Sun, Y., Liang, Y., Liu, Q., Shi, Y., Zhang, C. S., Zhang, C., Song, L., Zhang, P., Zhang, X., Li, X., Chen, T., Huang, H. Y., He, X., Wang, Y., Wu, Y. Q., Chen, S., Jiang, M., Chen, C.,… Lin, S. C. (2016). ULK1/2 Constitute a Bifurcate Node Controlling Glucose Metabolic Fluxes in Addition to Autophagy. Mol Cell, 62(3), 359–370. 10.1016/j.molcel.2016.04.009

Lin, X., Chen, Z., Yue, P., Averna, M. R., Ostlund, R. E., Jr., Watson, M. A., & Schonfeld, G. (2006). A targeted apoB38.9 mutation in mice is associated with reduced hepatic cholesterol synthesis and enhanced lipid peroxidation. Am J Physiol Gastrointest Liver Physiol, 290(6), G1170–1176. 10.1152/ajpgi.00402.2005

Lucas, S., Chen, G., Aras, S., & Wang, J. (2018). Serine catabolism is essential to maintain mitochondrial respiration in mammalian cells. Life Sci Alliance, 1(2), e201800036. 10.26508/lsa.201800036

Matyi, S., Jackson, J., Garrett, K., Deepa, S. S., & Unnikrishnan, A. (2018). The effect of different levels of dietary restriction on glucose homeostasis and metabolic memory. Geroscience, 40(2), 139–149. 10.1007/s11357-018-0011-5

Mitchell, S. J., Bernier, M., Mattison, J. A., Aon, M. A., Kaiser, T. A., Anson, R. M., Ikeno, Y., Anderson, R. M., Ingram, D. K., & de Cabo, R. (2019). Daily Fasting Improves Health and Survival in Male Mice Independent of Diet Composition and Calories. Cell Metab, 29(1), 221–228 e223. 10.1016/j.cmet.2018.08.011

Mteyrek, A., Filipski, E., Guettier, C., Okyar, A., & Levi, F. (2016). Clock gene Per2 as a controller of liver carcinogenesis. Oncotarget, 7(52), 85832–85847. 10.18632/oncotarget.11037

Neinast, M. D., Jang, C., Hui, S., Murashige, D. S., Chu, Q., Morscher, R. J., Li, X., Zhan, L., White, E., Anthony, T. G., Rabinowitz, J. D., & Arany, Z. (2019). Quantitative Analysis of the Whole-Body Metabolic Fate of Branched-Chain Amino Acids. Cell Metab, 29(2), 417–429 e414. 10.1016/j.cmet.2018.10.013

Ng, J. M., Azuma, K., Kelley, C., Pencek, R., Radikova, Z., Laymon, C., Price, J., Goodpaster, B. H., & Kelley, D. E. (2012). PET imaging reveals distinctive roles for different regional adipose tissue depots in systemic glucose metabolism in nonobese humans. Am J Physiol Endocrinol Metab, 303(9), E1134–1141. 10.1152/ajpendo.00282.2012

Nichenko, A. S., Sorensen, J. R., Southern, W. M., Qualls, A. E., Schifino, A. G., McFaline-Figueroa, J., Blum, J. E., Tehrani, K. F., Yin, H., Mortensen, L. J., Thalacker-Mercer, A. E., Greising, S. M., & Call, J. A. (2021). Lifelong Ulk1-Mediated Autophagy Deficiency in Muscle Induces Mitochondrial Dysfunction and Contractile Weakness. Int J Mol Sci, 22(4). 10.3390/ijms22041937

Okun, J. G., Rusu, P. M., Chan, A. Y., Wu, Y., Yap, Y. W., Sharkie, T., Schumacher, J., Schmidt, K. V., Roberts-Thomson, K. M., Russell, R. D., Zota, A., Hille, S., Jungmann, A., Maggi, L., Lee, Y., Bluher, M., Herzig, S., Keske, M. A., Heikenwalder, M.,… Rose, A. J. (2021). Liver alanine catabolism promotes skeletal muscle atrophy and hyperglycaemia in type 2 diabetes. Nat Metab, 3(3), 394–409. 10.1038/s42255-021-00369-9

Pak, H. H., Haws, S. A., Green, C. L., Koller, M., Lavarias, M. T., Richardson, N. E., Yang, S. E., Dumas, S. N., Sonsalla, M., Bray, L., Johnson, M., Barnes, S., Darley-Usmar, V., Zhang, J., Yen, C. E., Denu, J. M., & Lamming, D. W. (2021). Fasting drives the metabolic, molecular and geroprotective effects of a calorie-restricted diet in mice. Nat Metab, 3(10), 1327–1341. 10.1038/s42255-021-00466-9

Pareek, G., & Kundu, M. (2024). Physiological functions of ULK1/2. J Mol Biol, 436(15), 168472. 10.1016/j.jmb.2024.168472

Park, J. M., Lee, D. H., & Kim, D. H. (2023). Redefining the role of AMPK in autophagy and the energy stress response. Nat Commun, 14(1), 2994. 10.1038/s41467-023-38401-z

Parker, S. J., Amendola, C. R., Hollinshead, K. E. R., Yu, Q., Yamamoto, K., Encarnacion-Rosado, J., Rose, R. E., LaRue, M. M., Sohn, A. S. W., Biancur, D. E., Paulo, J. A., Gygi, S. P., Jones, D. R., Wang, H., Philips, M. R., Bar-Sagi, D., Mancias, J. D., & Kimmelman, A. C. (2020). Selective Alanine Transporter Utilization Creates a Targetable Metabolic Niche in Pancreatic Cancer. Cancer Discov, 10(7), 1018–1037. 10.1158/2159-8290.CD-19-0959

Paulusma, C. C., Lamers, W. H., Broer, S., & van de Graaf, S. F. J. (2022). Amino acid metabolism, transport and signalling in the liver revisited. Biochem Pharmacol, 201, 115074. 10.1016/j.bcp.2022.115074

Petersen, K. F., Dufour, S., Cline, G. W., & Shulman, G. I. (2019). Regulation of hepatic mitochondrial oxidation by glucose-alanine cycling during starvation in humans. J Clin Invest, 129(11), 4671–4675. 10.1172/JCI129913

Reinke, H., & Asher, G. (2016). Circadian Clock Control of Liver Metabolic Functions. Gastroenterology, 150(3), 574–580. 10.1053/j.gastro.2015.11.043

Remesy, C., Moundras, C., Morand, C., & Demigne, C. (1997). Glutamine or glutamate release by the liver constitutes a major mechanism for nitrogen salvage. Am J Physiol, 272(2 Pt 1), G257–264. 10.1152/ajpgi.1997.272.2.G257

Rhoads, T. W., Clark, J. P., Gustafson, G. E., Miller, K. N., Conklin, M. W., DeMuth, T. M., Berres, M. E., Eliceiri, K. W., Vaughan, L. K., Lary, C. W., Beasley, T. M., Colman, R. J., & Anderson, R. M. (2020). Molecular and Functional Networks Linked to Sarcopenia Prevention by Caloric Restriction in Rhesus Monkeys. Cell Syst, 10(2), 156–168 e155. 10.1016/j.cels.2019.12.002

Riera, C. E., & Dillin, A. (2015). Tipping the metabolic scales towards increased longevity in mammals. Nat Cell Biol, 17(3), 196–203. 10.1038/ncb3107

Rohart, F., Gautier, B., Singh, A., & Le Cao, K. A. (2017). mixOmics: An R package for ‘omics feature selection and multiple data integration. PLoS Comput Biol, 13(11), e1005752. 10.1371/journal.pcbi.1005752

Sanford, J. D., Franklin, D., Grois, G. A., Jin, A., & Zhang, Y. (2023). Carnitine o-octanoyltransferase is a p53 target that promotes oxidative metabolism and cell survival following nutrient starvation. J Biol Chem, 299(7), 104908. 10.1016/j.jbc.2023.104908

Sarabhai, T., & Roden, M. (2019). Hungry for your alanine: when liver depends on muscle proteolysis. J Clin Invest, 129(11), 4563–4566. 10.1172/JCI131931

Shao, D., Villet, O., Zhang, Z., Choi, S. W., Yan, J., Ritterhoff, J., Gu, H., Djukovic, D., Christodoulou, D., Kolwicz, S. C., Jr., Raftery, D., & Tian, R. (2018). Glucose promotes cell growth by suppressing branched-chain amino acid degradation. Nat Commun, 9(1), 2935. 10.1038/s41467-018-05362-7

Sharma, N., Sequea, D. A., Castorena, C. M., Arias, E. B., Qi, N. R., & Cartee, G. D. (2014). Heterogeneous effects of calorie restriction on in vivo glucose uptake and insulin signaling of individual rat skeletal muscles. PLoS One, 8(6), e65118. 10.1371/journal.pone.0065118

Sousa, C. M., Biancur, D. E., Wang, X., Halbrook, C. J., Sherman, M. H., Zhang, L., Kremer, D., Hwang, R. F., Witkiewicz, A. K., Ying, H., Asara, J. M., Evans, R. M., Cantley, L. C., Lyssiotis, C. A., & Kimmelman, A. C. (2016). Pancreatic stellate cells support tumour metabolism through autophagic alanine secretion. Nature, 536(7617), 479–483. 10.1038/nature19084

Specht, K. S., Kant, S., Addington, A. K., McMillan, R. P., Hulver, M. W., Learnard, H., Campbell, M., Donnelly, S. R., Caliz, A. D., Pei, Y., Reif, M. M., Bond, J. M., DeMarco, A., Craige, B., Keaney, J. F., Jr., & Craige, S. M. (2021). Nox4 mediates skeletal muscle metabolic responses to exercise. Mol Metab, 45, 101160. 10.1016/j.molmet.2020.101160

Teong, X. T., Liu, K., Vincent, A. D., Bensalem, J., Liu, B., Hattersley, K. J., Zhao, L., Feinle-Bisset, C., Sargeant, T. J., Wittert, G. A., Hutchison, A. T., & Heilbronn, L. K. (2023). Intermittent fasting plus early time-restricted eating versus calorie restriction and standard care in adults at risk of type 2 diabetes: a randomized controlled trial. Nat Med, 29(4), 963–972. 10.1038/s41591-023-02287-7

Veldhorst, M. A., Westerterp-Plantenga, M. S., & Westerterp, K. R. (2009). Gluconeogenesis and energy expenditure after a high-protein, carbohydrate-free diet. Am J Clin Nutr, 90(3), 519–526. 10.3945/ajcn.2009.27834

Wang, H., Arias, E. B., Yu, C. S., Verkerke, A. R. P., & Cartee, G. D. (2017). Effects of Calorie Restriction and Fiber Type on Glucose Uptake and Abundance of Electron Transport Chain and Oxidative Phosphorylation Proteins in Single Fibers from Old Rats. J Gerontol A Biol Sci Med Sci, 72(12), 1638–1646. 10.1093/gerona/glx099

Wang, T. J., Larson, M. G., Vasan, R. S., Cheng, S., Rhee, E. P., McCabe, E., Lewis, G. D., Fox, C. S., Jacques, P. F., Fernandez, C., O’Donnell, C. J., Carr, S. A., Mootha, V. K., Florez, J. C., Souza, A., Melander, O., Clish, C. B., & Gerszten, R. E. (2011). Metabolite profiles and the risk of developing diabetes. Nat Med, 17(4), 448–453. 10.1038/nm.2307

Wang, X., Dong, L. Y., Gai, Q. J., Ai, W. L., Wu, Y., Xiao, W. C., Zhang, J., & An, W. (2020). Lack of Augmenter of Liver Regeneration Disrupts Cholesterol Homeostasis of Liver in Mice by Inhibiting the AMPK Pathway. Hepatol Commun, 4(8), 1149–1167. 10.1002/hep4.1532

Wu, Y., Xiao, H., Zhang, H., Pan, A., Shen, J., Sun, J., Liang, Z., & Pi, J. (2023). Quasi-Targeted Metabolomics Approach Reveal the Metabolite Differences of Three Poultry Eggs. Foods, 12(14). 10.3390/foods12142765

Yang, L., Garcia Canaveras, J. C., Chen, Z., Wang, L., Liang, L., Jang, C., Mayr, J. A., Zhang, Z., Ghergurovich, J. M., Zhan, L., Joshi, S., Hu, Z., McReynolds, M. R., Su, X., White, E., Morscher, R. J., & Rabinowitz, J. D. (2020). Serine Catabolism Feeds NADH when Respiration Is Impaired. Cell Metab, 31(4), 809–821 e806. 10.1016/j.cmet.2020.02.017

Yoon, I., Nam, M., Kim, H. K., Moon, H. S., Kim, S., Jang, J., Song, J. A., Jeong, S. J., Kim, S. B., Cho, S., Kim, Y., Lee, J., Yang, W. S., Yoo, H. C., Kim, K., Kim, M. S., Yang, A., Cho, K., Park, H. S.,… Kim, S. (2020). Glucose-dependent control of leucine metabolism by leucyl-tRNA synthetase 1. Science, 367(6474), 205–210. 10.1126/science.aau2753

Zhou, Q., Yu, L., Cook, J. R., Qiang, L., & Sun, L. (2023). Deciphering the decline of metabolic elasticity in aging and obesity. Cell Metab, 35(9), 1661–1671 e1666. 10.1016/j.cmet.2023.08.001

