## Supplementary figures and images for "Ulk1(S555) inhibition alters nutrient stress response by prioritizing amino acid metabolism"

### supplemental figures

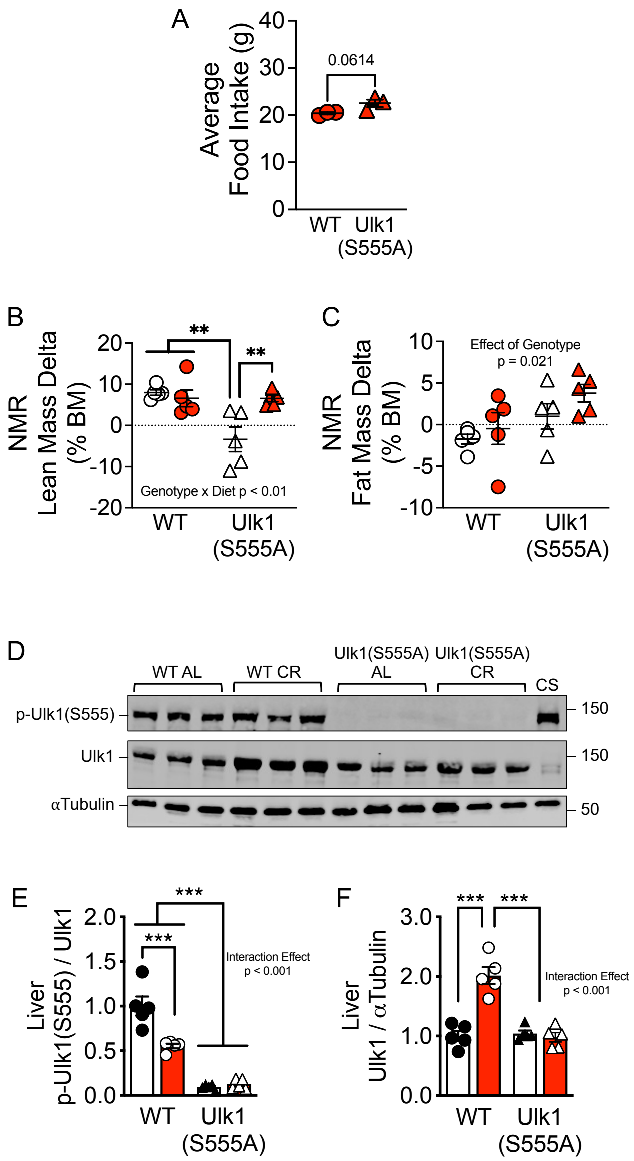

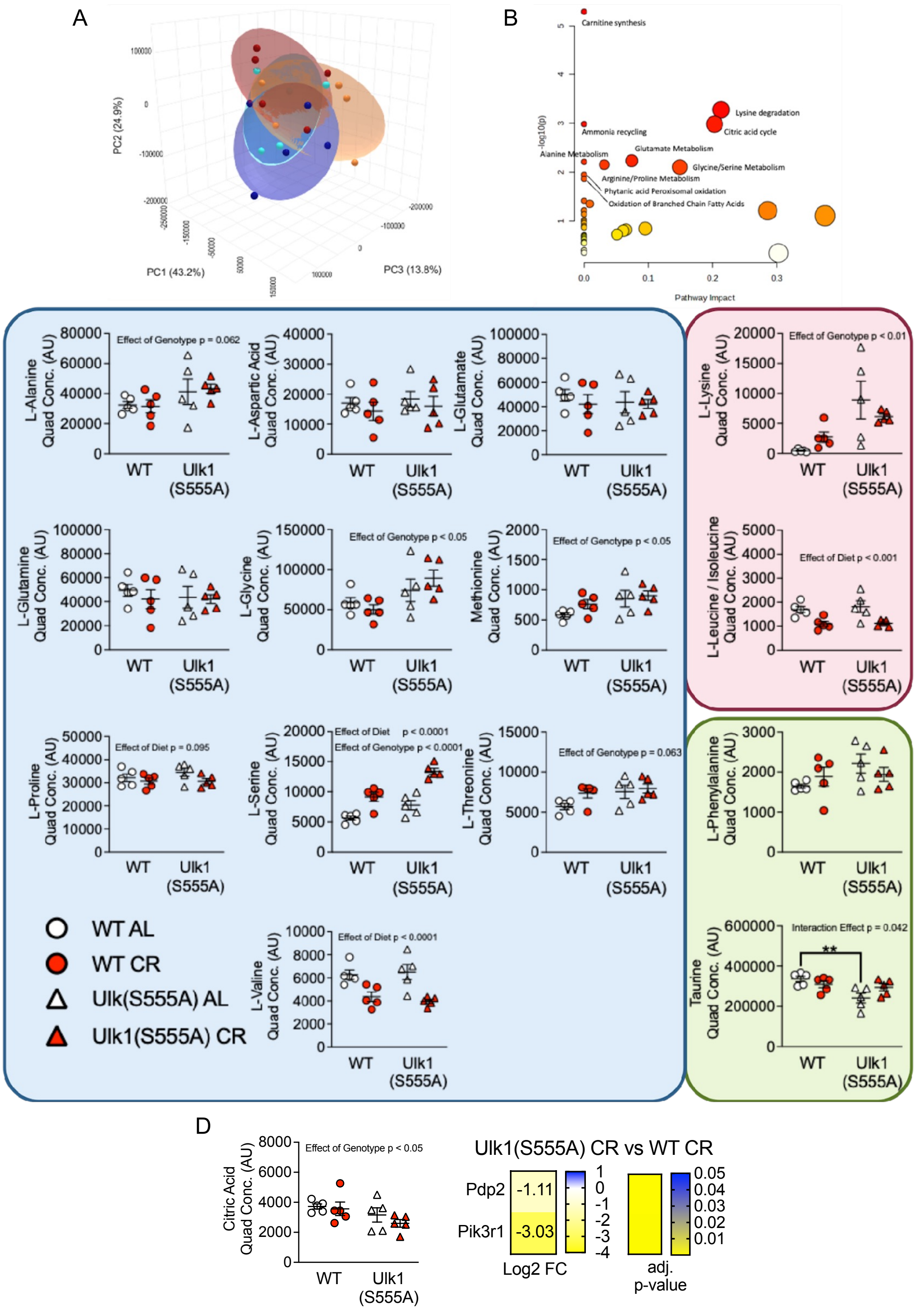

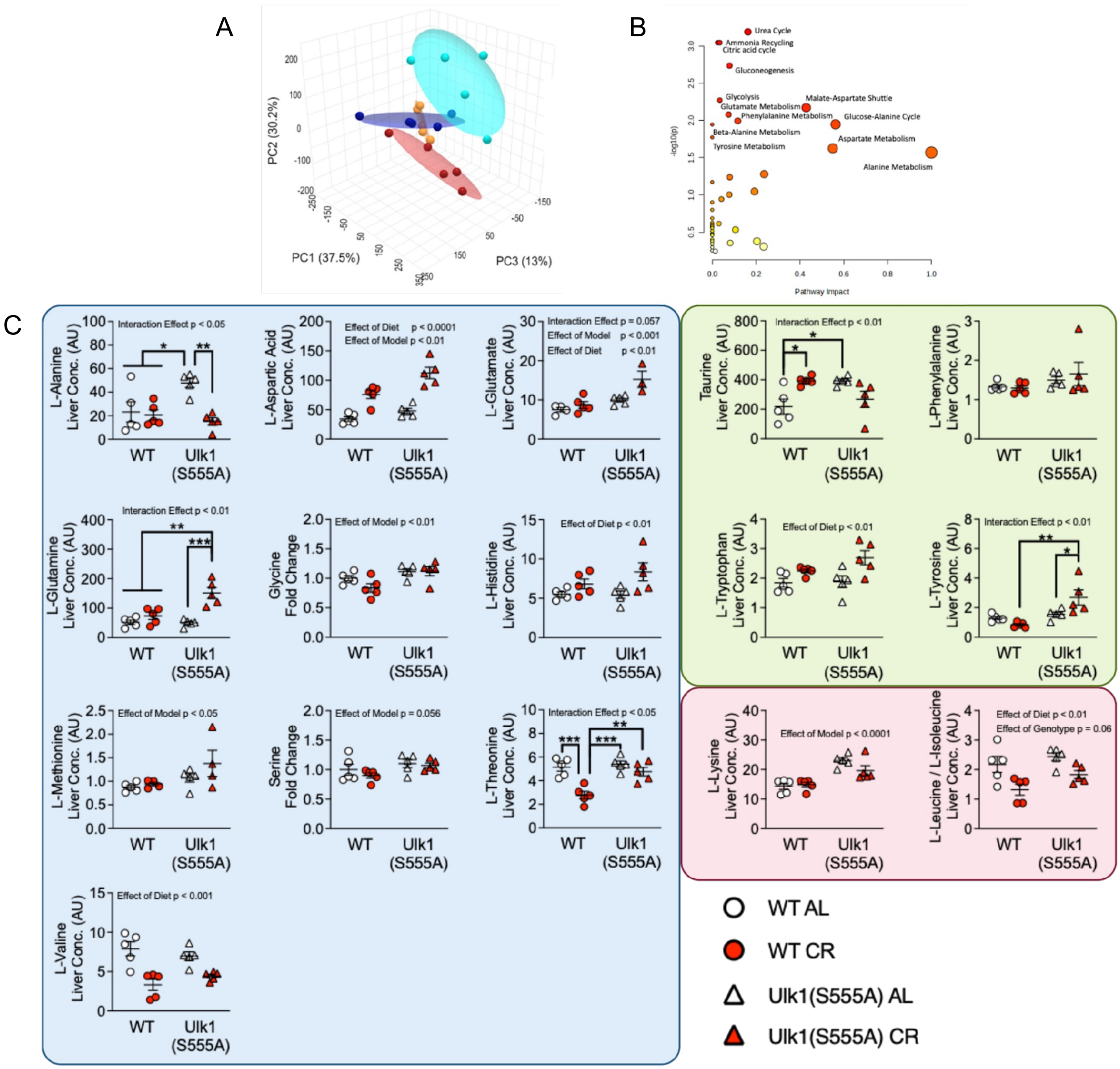

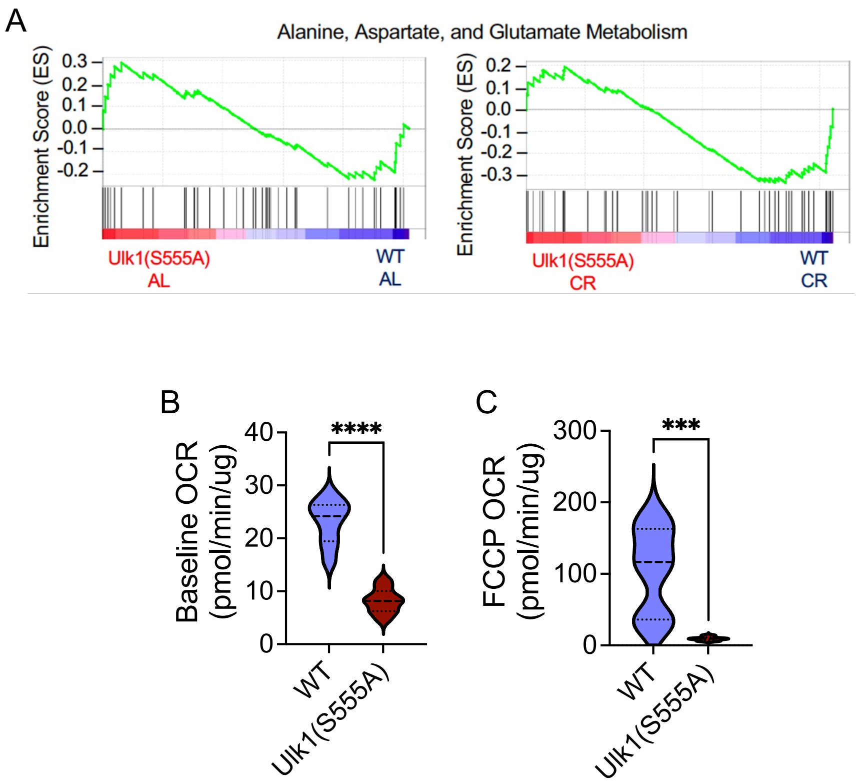

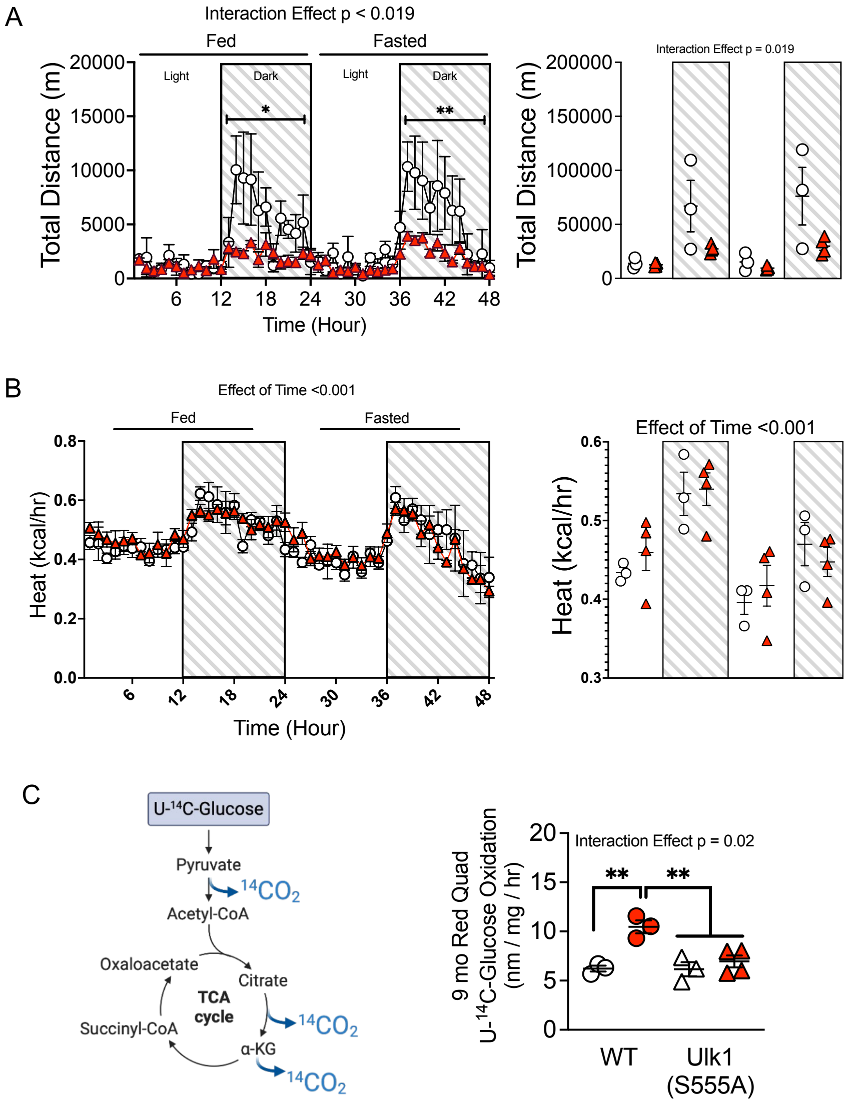

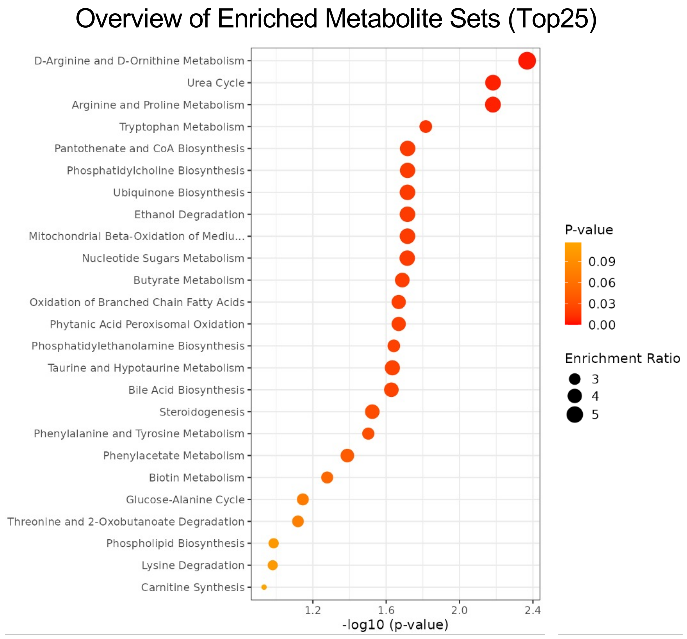

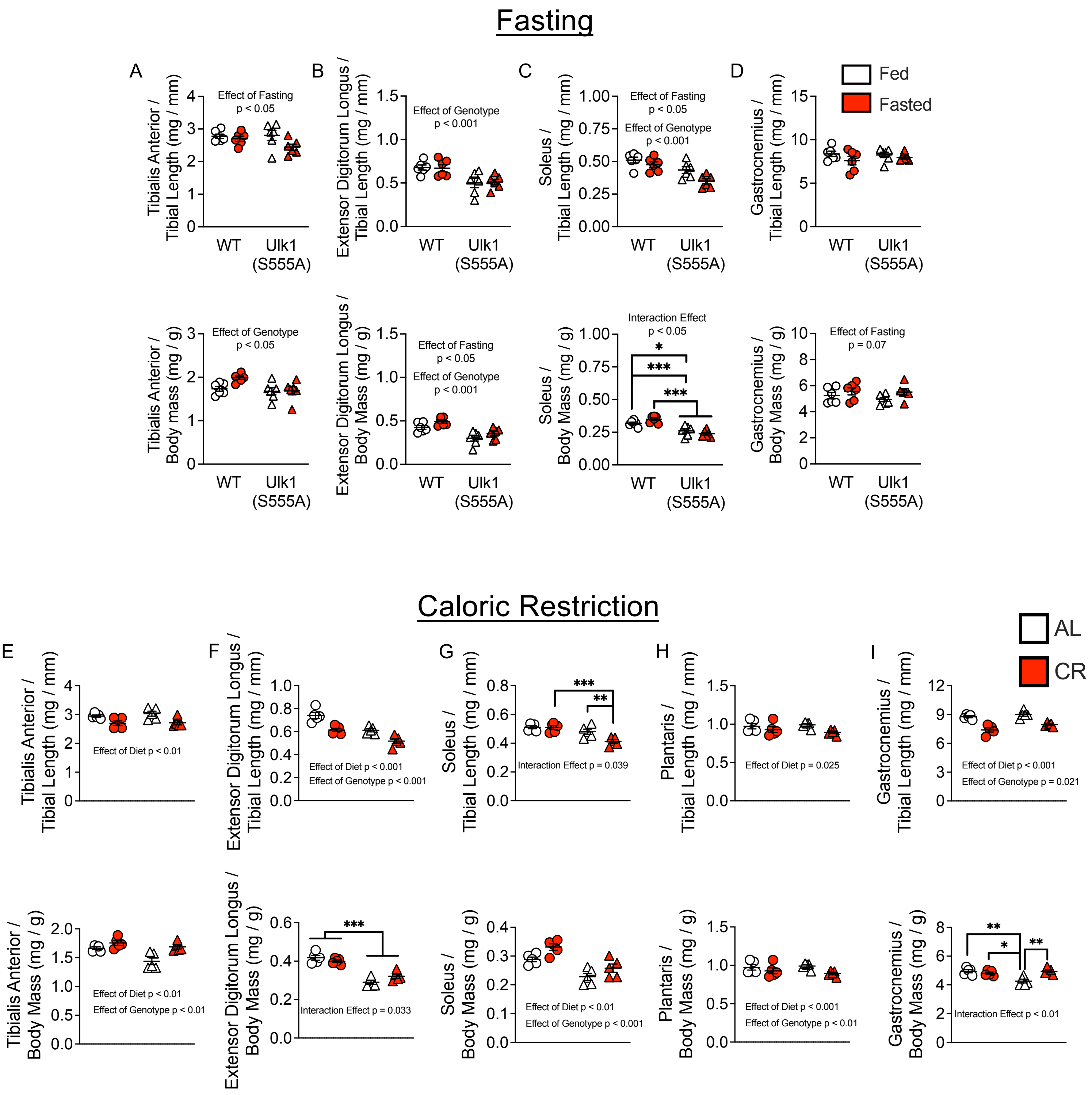

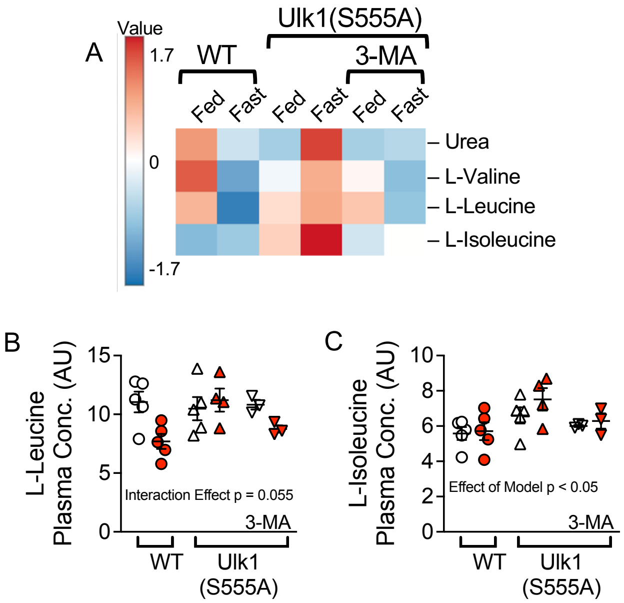
